# Conditions for successful nitrogen removal from source-separated urine by partial nitritation/anammox

**DOI:** 10.1101/2023.12.22.573048

**Authors:** Valentin Faust, Philipp Markus, Sarina Schielke-Jenni, Marijn J. Timmer, Jolien De Paepe, Ramon Ganigué, Siegfried E. Vlaeminck, Kai M. Udert

**Affiliations:** Eawag, Swiss Federal Institute of Aquatic Science and Technology, 8600 Dübendorf, Switzerland; ETH Zürich, Institute of Environmental Engineering, 8093 Zürich, Switzerland; HFS Aqua AG, 8400 Winterthur, Switzerland; AWEL Department for Waste, Water, Energy, and Air, 8090 Zürich, Switzerland; Research Group of Sustainable Energy, Air and Water Technology, Department of Bioscience Engineering, Faculty of Science, University of Antwerp, 2020 Antwerpen, Belgium; Center for Advanced Process Technology for Urban Resource Recovery (CAPTURE), Frieda Saeysstraat 1, 9052 Gent, Belgium; Center for Microbial Ecology and Technology, Department of Biotechnology, Faculty of Bioscience Engineering, Ghent University, Coupure Links 653, 9000 Gent, Belgium

**Keywords:** Source separation, Nitrogen removal, Decentralized treatment, Regenerative life support systems

## Abstract

Partial nitritation/anammox (PN/A) of source-separated urine is less energy-intensive and potentially cheaper and more environmentally friendly than conventional nitrogen removal from mixed sewage. However, PN/A of undiluted source-separated urine has not yet been established. In this study, the feasibility of PN/A for source-separated urine (total nitrogen ≈ 2 to 3 g-N L^-1^). To evaluate the influence of different factors, one- and two-stage configurations were operated using different influents, i.e. source-separated urine, synthetic urine, and urine with additional divalent cations. While partial nitritation was successfully achieved in both configurations with digester supernatant and urine, anammox activity was lost shortly after switching from digester supernatant to the urine influents. Toxic organic compounds or pharmaceuticals and the high monovalent to divalent cation ratio were suspected as causes of anammox failure, but were ruled out due to the different reactor configurations and influent compositions tested. Other suspected factors such as COD/N ratio, phosphate and sulfate inhibition, nitrogen compound inhibition, metal inhibition, pH and dissolved oxygen were also systematically excluded. Instead, the high salt concentration in urine compared to the digester supernatant most likely caused the reactor to fail due to the disintegration of large flocs, and the resulting challenge of biomass retention. The shortcomings of the floccular sludge system were overcome by using biofilm carriers, resulting in successful PN/A. This hybrid system ran for 140 days with nitrogen removal rates of up to 1000 mg-N L^-1^ d^-1^ with an average of 410 ± 220 mg-N L^-1^ d^-1^, and a nitrogen removal efficiency of 93 ± 3% at 30°C.

## 1 Introduction

Urine contributes about 80% of the nitrogen in domestic wastewater, but only to around 14% of the organics expressed as chemical oxygen demand (COD) (Larsen and Gujer, 1996). The total ammoniacal nitrogen (TAN = NH_3_-N + NH_4_ ^+^-N) concentration in stored urine, i.e. after urea hydrolysis, can be as high as 8 g-N L^-1^, typically at a COD/N ratio of around 1 (Udert et al., 2006). However, due to mixing with some flushing water, the TAN concentration of source-separated urine is typically between 2 and 4 g-N L^-1^ (Fumasoli, 2016). Decentralized nitrogen treatment of source-separated urine is potentially cheaper and less energy-intensive than for wastewater, where the nitrogen is already highly diluted (Maurer et al., 2006; Wilsenach and Loosdrecht, 2006). If about 60-75% of the urine is treated on-site, the nitrogen treatment step in wastewater treatment plants would no longer be necessary, as all the remaining nitrogen would be used for assimilation (Wilsenach and van Loosdrecht, 2003), which could also yield a lower greenhouse gas emission footprint. Typically, urine treatment is associated with fertiliser production by recovering the nitrogen as, for example, struvite or ammonium nitrate (Larsen ^e^t al., 202^1^). However, nitrogen removal as nitrogen gas (N_2_) rather than recovery may be more economically and energetically attractive due to the well-established fertiliser production pathway via the Haber-Bosch process (Spiller et al., 2022). In the special case of space missions, converting nitrogen to N_2_ could be used to counteract gas losses and provide atmospheric pressure in regenerative biological life support systems, or provide a nitrogen source for food production based on biological nitrogen fixation, uncoupled from potentially undesired residuals in the urine matrix (Timmer et al., 2023).

The conventional process for biological nitrogen removal from wastewater combines autotrophic nitrification and heterotrophic denitrification, consuming at least around 4 g-COD per g-N removed (Tchobanoglous et al., 2014). Hence, without dosing additional organics (COD), traditional nitrification/denitrification is not feasible with urine (Schielke-Jenni et al., 2015). In contrast, nitrogen removal via partial nitritation and anaerobic ammonium oxidation (anammox), abbreviated as PN/A, is fully autotrophic and does not require any biodegradable organic substrate, although some contribution of heterotrophic denitrification cannot be ruled out in real systems (Schielke-Jenni et al., 2015). In addition, PN/A is economically advantageous as it reduces the oxygen requirement by up to 60% compared to conventional nitrification/denitrification (Siegrist et al., 2008). PN/A can either be operated in one-stage or two-stage configurations. One-stage reactors are more common, partly because of their smaller footprint and lower capital investment (Lackner et al., 2014). Systems are operated either with floccular sludge, with biofilm, growing on carrier material, or as a hybrid system. Hybrid systems, e.g. integrated fixed-film activated sludge systems, have the potential to offer better performance than pure biofilm-based systems (Wang et al., 2022). However, the addition of biofilm carriers will increase the costs and may increase the energy required for aeration and mixing (Nelson et al., 2021).

The PN/A application on sewage sludge reject water, termed sidestream nitrogen removal, is a mature technology with over 100 full-scale installations worldwide (Lackner et al., 2014). However, the composition of urine is very different from the usual influent of high-strength nitrogen PN/A processes, i.e., digester supernatant, due to the high concentration of organics and the high salinity. Stored urine has a COD to nitrogen ratio of about 1 g-COD g-N^-1^ (Faust et al., 2023b), which is higher than for digester supernatant, at about 0.4 g-COD g-N^-1^ (Udert et al., 2008). Organic substances can potentially be directly toxic to the bacteria involved in the PN/A process (Gomez et al., 2000; Yang et al., 2013). Moreover, two-thirds of pharmaceuticals such as antibiotics are excreted in the urine (Lienert et al., 2007), where they are present in higher concentrations than in diluted sewage. These pharmaceuticals can be toxic to bacteria (Fernández et al., 2009). In addition, organic substances promote the growth of heterotrophic bacteria, which compete with aerobic ammonia-oxidizing bacteria (AerAOB) for oxygen and with anammox bacteria (AnAOB) for nitrite (Chamchoi et al., 2008). However, Jenni et al. (2014) showed that AnAOB can coexist with HET at least up to influent COD to N ratios of 1.4 g-COD g-N^-1^ provided a high sludge retention time (SRT) is maintained (Jenni et al., 2014).

Stored source-separated urine has high salinities of 10 to 30 mS cm^-1^, which can directly inhibit bacteria. (Moussa et al., 2006). AnAOB have been shown to be sensitive to salinity, but can also adapt to high salt concentrations if the salinity is gradually increased (Kartal et al., 2006). Faust et al. (2023b) showed that an increase in salinity during urine nitrification strongly reduces the activity of AerAOB and nitrite-oxidizing bacteria (NOB). As the decrease in activity was greater for AerAOB than for NOB, the increase in salinity may be problematic for partial nitritation. In addition to the high salinity, urine has a high ratio of monovalent to divalent cations, due to high concentrations of sodium, ammonium and potassium compared to magnesium and calcium, resulting in a ratio of 400 mol mol^-1^ compared to 20 mol mol^-1^ in the digester supernatant (Jenni, 2015). The high ratio of monovalent to divalent cations, can affect floc formation and lead to destabilization of flocs (Dapena-Mora et al., 2010), posing a challenge to sludge retention. The divalent cation bridging theory states that divalent cations can bridge negatively charged functional groups of extracellular polymeric substances. This bridging helps aggregate and stabilise the biopolymer and bacterial matrix, promoting bioflocculation (Higgins et al., 2004). Sobeck and Higgins (2002) showed that the addition of sodium resulted in a deterioration of floc properties (sludge volume index, floc strength, capillary suction time), whereas the addition of either calcium or magnesium resulted in an improvement of floc properties.

The operation of a one-stage PN/A process with highly diluted urine (TAN ≈ 0.6 g-N L^-1^) in a floccular sludge system was successfully tested in two previous studies but was prone to process failures due to a loss of AnAOB activity (Bürgmann et al., 2011; Udert et al., 2008). Only with a biofilm system, i.e., a membrane-aerated biofilm reactor (MABR), were Timmer et al. (2023) able to achieve stable nitrogen removal rates (NRR) as gaseous nitrogen (N^2^) of up to 1 g-N L^-1^ d^-1^ and nitrogen removal efficiencies (NRE) of 80-95% on diluted urine (total nitrogen ≈ 0.7 g-N L^-1^). In addition, the MABR was successfully tested for a short period of 50 days on source-separated urine with nitrogen concentrations of 2 to 5.5 g-N L^-1^. Instead of hydrolyzed urine (pH ≈ 9), stabilized urine (pH ≈ 11) was fed to the reactor, i.e. base was added to the fresh urine to prevent urea hydrolysis during storage, which also adds alkalinity to the urine. Pourbavarsad et al. (2021) also used a MABR to treat source-separated urine with nitrogen concentrations of 2 to 5.5 g-N L^-1^, but NRR of only 85 mg-N L^-1^ d^-1^ and NRE of 0-40% were reached. Partial nitritation of urine has been successfully achieved at various pH values ranging from 4.5 to 8.5 (Faust et al., 2022; Faust et al., 2023b; Pourbavarsad et al., 2022), but the combination with anammox in a two-step configuration has yet to be tested.

Until now, PN/A with source-separated urine could only be achieved in a floccular sludge system with highly diluted urine or in an MABR using undiluted pre-alkalinized urine. So far, the systems with floccular sludge systems were prone to failure. A robust, simple, and inexpensive process is essential for decentralised treatment. The aim of this study was to test the feasibility of PN/A for source-separated urine (TAN ≈ 2 to 3 g-N L^-1^) using a floccular system. The main focus was on one-stage PN/A configurations. However, for the first time, a two-stage system was also investigated with urine, as this allowed the nitritation and anammox processes to be studied separately. Additional attention was given to the effects of potentially toxic compounds, e.g. pharmaceuticals, and the ratio of monovalent to divalent cations. Finally, biofilm carriers were added to the floccular sludge system to see if a hybrid system would improve PN/A treatment.

## 2 Materials and methods

### 2.1 Reactor operation and influent composition

Five experiments were carried out with different reactor configurations and influent compositions (**Table 1**).

**Table 1:**
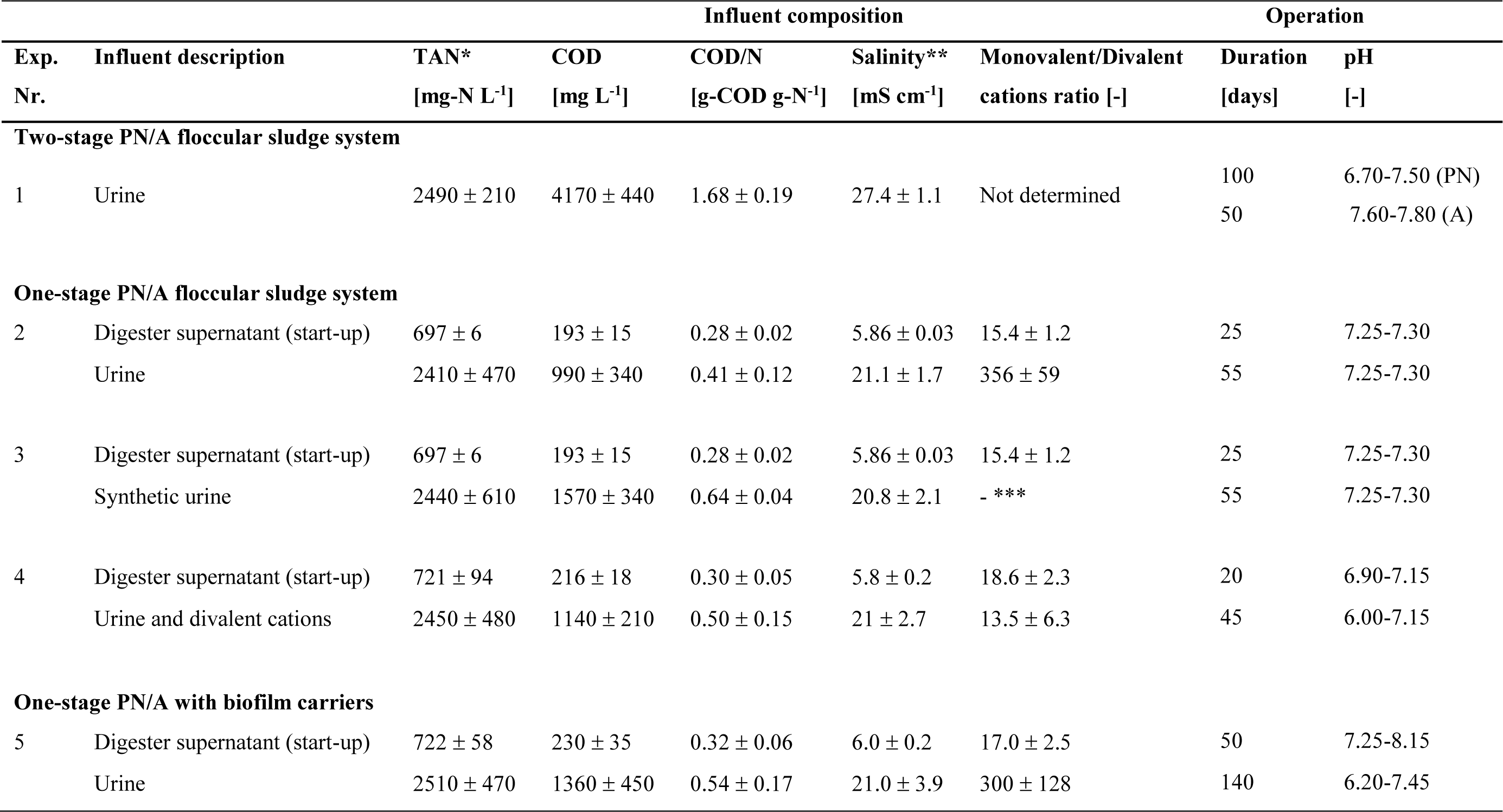
Influent compositions, experiment duration and operational pH for all five experiments. In experiment Nr. 2 to Nr. 5, the reactors were first fed with digester supernatant before the influent was changed. See **Supplementary Information (SI) 1** for more information about the influent composition. *Total ammoniacal nitrogen (TAN = NH_3_-N + NH^4+^-N). **Electrical conductivity was used as a proxy for salinity. *** No divalent cations were added to the synthetic urine.

#### 2.1.1 Two-stage partial nitritation/anammox with floccular sludge

A two-stage PN/A configuration using floccular sludge was investigated to assess the effects on nitritation and nitrogen removal with anammox separately (**Fig 1**). The partial nitritation reactor consisted of an aerated 7-L continuous-flow stirred reactor (CSTR) without additional sludge retention, i.e. the SRT was equal to the hydraulic retention time (HRT). The partial nitritation reactor was inoculated with suspended sludge from the nitrification stage of the wastewater treatment plant at Eawag (Dübendorf, Switzerland). Stored source-separated urine, from waterless urinals and source-separating toilets was used as influent (experiment Nr. 1 in **Table 1**). The pH in the reactor was controlled between 6.7 and 7.5 using the influent (stored urine pH ≈ 9). The temperature was controlled at 30°C and constant aeration was provided at 1 L min^-1^.

**Fig 1:**
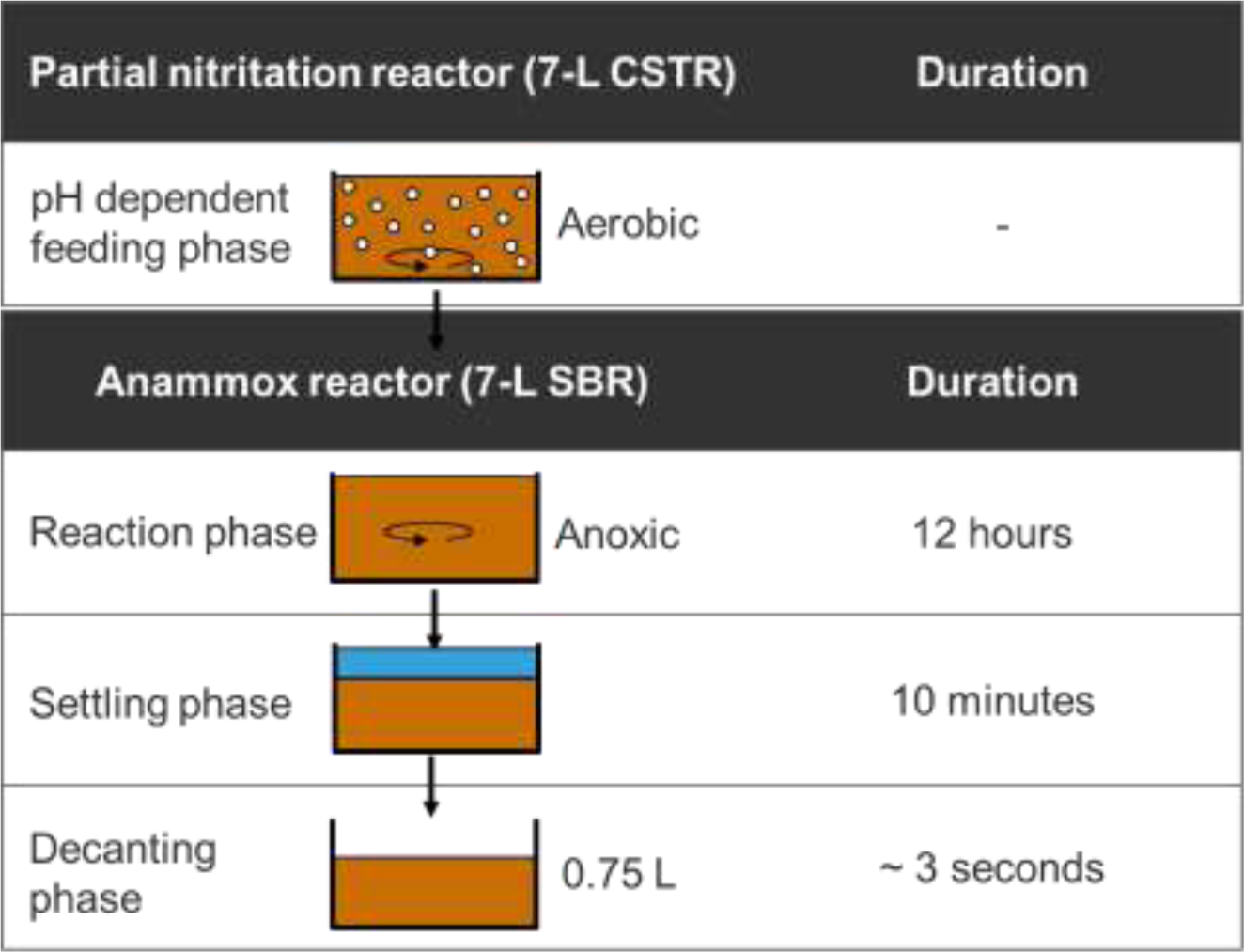
Set-up of two-stage reactor. Continuous-flow stirred reactor (CSTR) for partial nitritation followed by a sequencing batch reactor (SBR) for anammox consisting of (i) reaction phase with anoxic stirring, (ii) sedimentation, and (iii) withdrawal.

After the nitritation reactor had been running for about 55 days, the anoxic reactor for anammox was started using suspended biomass from a PN/A reactor treating sludge digester supernatant (WWTP Limmattal, Dietikon, Switzerland) as inoculum. The anoxic reactor consisted of a 7-L sequencing batch reactor (SBR). The SBR consisted of a stirred, anoxic phase, a settling phase of 10 min and a short withdrawal phase. Initially, the length of the anoxic phase was time-controlled, i.e. 12 h, and the exchange volume per cycle was 0.75 L per cycle. From day 10 onwards, the length of the anoxic phase was extended due to the slow AnAOB activity. To avoid ammonia toxicity and ensure high NRE, the ammonium concentration at the end of each cycle was kept below 50 mg-N L^-1^ by adding additional nitrite. The pH was controlled at 7.6 to 7.8 by adding hydrochloric acid, and the temperature was maintained at 30°C.

#### 2.1.2 One-stage partial nitritation/anammox with floccular sludge

One-stage PN/A configurations were tested using floccular sludge. To investigate the effect of bulk organics, pharmaceuticals, salt concentrations, and the type of cations, three experiments were conducted with different influent compositions (experiment Nr. 2, 3 and 4 in **Table 1**). The three 13-L reactors were operated in fed-batch mode. Fed-batch mode was chosen over a sequential batch reactor (SBR) to avoid large pH fluctuations (Faust et al., 2023a). A cycle always consisted of four different steps: (i) feeding phase, (ii) stirring phase, (iii) settling phase, and (iv) decanting phase (**Fig 2**). (i) The first step consisted of a pH-dependent feeding with alternating aerated and anoxic phases until the maximum fill level of 13 L was reached. Due to the proton released during ammonia oxidation, the pH decreased. When the lower pH setpoint was reached, the influent was switched on, which increased the pH due to the higher pH and alkalinity in the influent (stored urine pH ≈ 9). In experiment Nr. 2 and 3, the pH was kept constant between 7.25 and 7.30, and in experiment Nr. 4 the pH was adjusted to optimize the process (**Table 1**). In principle, lower pH set-points were tested to reduce the inhibition of AnAOB by free ammonia, while providing sufficient substrate for AerAOB. The aeration intensity in all reactors was manually adjusted as a function of the DO in the reactor to ensure values around 0.1-0.2 mg-O_2_ L^-1^ during the aerobic phase. The duration of the aerated phase was manually adjusted according to the nitrite production by AerAOB. The anoxic phase was manually adjusted as a function of the nitrite degradation until no nitrite was present in the reactor, i.e. if nitrite was still present, the proportion of the feeding phase without aeration was increased. Adjustments were made at least once a week, except during holidays. (ii) Additional anoxic stirring was included prior to sedimentation to remove any remaining nitrite. (iii) Stirring was then stopped to allow the sludge to settle, and finally (iv) 1 L was withdrawn from the reactor. The temperature in the reactors was controlled at 30°C. For more information on the reactor set up, see **SI 2**.

**Fig 2:**
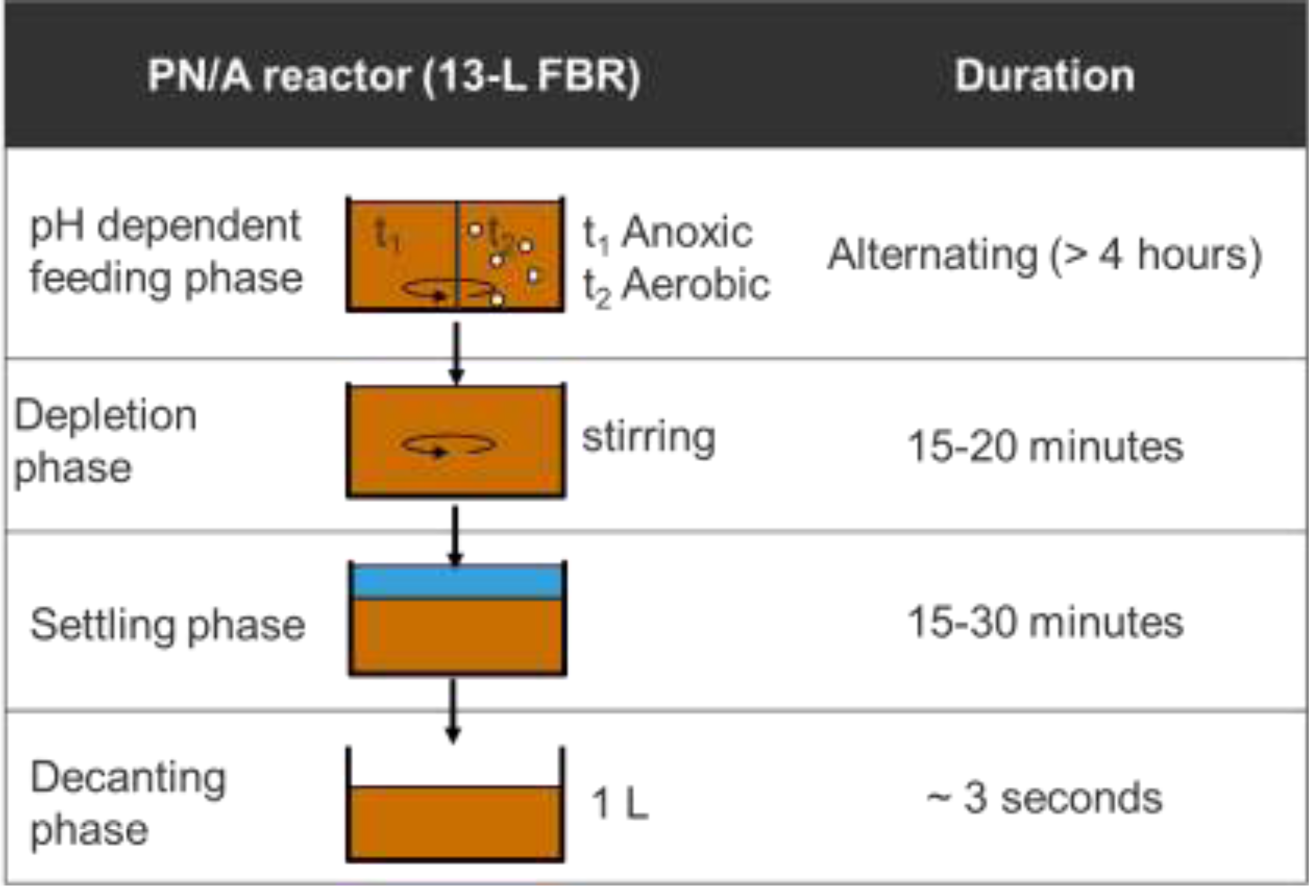
Set-up of one-stage reactor. Fed-batch reactor (FBR) consisting of (i) pH-dependent feeding with alternating aerobic and anoxic conditions, (ii) anoxic stirring, (iii) sedimentation, and (iv) withdrawal.

All three reactors were inoculated with 13 L of active PN/A floccular sludge taken from an SBR treating the digester supernatant of the sewage sludge treatment plant ZSA Pfannenstiel (Oetwil, Switzerland). For the start-up phase, each reactor was initially operated with digester supernatant from ZSA Pfannenstiel (Oetwil, Switzerland) for 3 to 4 weeks before switching to the specific influent (**Table 1**). In experiment Nr. 2, the reactor was operated with source-separated urine without any modifications and used as a reference reactor. In experiment Nr. 3, the reactor was operated with synthetic urine (recipe in **SI 1**) to test whether the absence of potentially toxic organic or pharmaceuticals would result in a successful PN/A process. The synthetic urine contained all major inorganic salts. In order to maintain the COD to N ratio of urine, acetate, which is the major organic compound in urine (Heusser et al., 2023), was added to the synthetic urine. In experiment Nr. 4, the reactor was operated with source-separated urine and additional divalent cations to determine if this stabilizes the process. The ratio of monovalent to divalent cations corresponded to the ratio found in digester supernatant.

#### 2.1.3 One-stage partial nitritation/anammox with floccular sludge and biofilm carriers

An additional reactor was operated to test whether PN/A on source-separated urine could be improved when using a combination of biofilm carriers (FLUOPUR, HFS Aqua) and floccular sludge, i.e a hybrid system. The carriers were retained with a sieve. The reactor was inoculated with biofilm carriers and floccular sludge from a PN/A reactor treating the digester supernatant at Langmatt (Wildegg, Switzerland). The reactor was operated as described in **Section 2.1.2** and in **Table 1** (experiment Nr. 5). The pH was adjusted to optimize the process.

### 2.2 Chemical and physical analyses

To analyse the dissolved compounds, samples were filtered through 0.45 µm GF/PET filters (Chromafil, Macherey-Nagel). Ion chromatography (881 compact IC pro, Metrohm) was used to measure cations (potassium, ammonium, and sodium) and anions (chloride, nitrite, phosphate, nitrate, and sulfate). Nitrite and nitrate concentrations were additionally checked using semi-quantitative colorimetric strips (110007 and 110020 MQuant, Merck). The acid-base equilibria of HNO_2_ and NO_2_ ^-^, and NH_4_ ^+^ and NH_3_ were calculated according to Crittenden et al. (2012) using the dissociation constants of Anthonisen et al. (1976) and corrected for ionic strength according to Davies (1967) (see **SI 3** and **SI 4**). Total suspended solids and volatile suspended solids were measured according to the APHA (2012) standard protocol. Dissolved COD was measured with a spectrophotometer (DR 2800, Hach Lange) using photometric cuvette tests (LCK114, Hach Lange). Electrical conductivity was measured with a standard conductivity cell (LF340, WTW) and corrected for the temperature at 25°C. The pH and the temperature were measured with glass electrodes (Orbisint CPS11D, Endress+Hauser), which were calibrated weekly. DO was measured with optical oxygen sensors (Oxymax COS61D and Memosens COS81D, Endress+Hauser). The airflow was controlled using an airflow controller (red-y smart controller GSC, Voegtlin).

### 2.3 Molecular analyses of the biomass

Biomass was sampled from Reactor 2 (floccular sludge) and Reactor 5 (floccular sludge and biofilm carriers), centrifuged, and stored at -20°C before further processing. DNA was extracted using the PowerSoil extraction kit (Qiagen, Hilden, Germany) according to the manufacturer’s protocol. DNA extracts were sent to Novogene (Beijing, China) for 16S rRNA gene-based amplicon sequencing, library preparation and sequencing on a Novaseq platform. The universal primer pair 341F / 806R was used, targeting the V3-V4 hypervariable region of bacterial 16S rRNA gene sequences (Klindworth et al., 2013).

## 3 Results

### 3.1 Two-stage partial nitritation/anammox with floccular sludge

While partial nitritation was successfully achieved in a two-stage configuration, it was not possible to successfully establish an anammox process using the partially-nitrified urine. In the partial nitritation reactor, ammonia oxidation rates of 315 ± 130 mg-N L^-1^ d^-1^ and low nitrate production were observed (**Fig 3**). The ratio of nitrite to ammonium in the effluent of the first reactor was 0.9 ± 0.05 g-N g-N^-1^ before base addition excluding the start-up period of 10 days. With the addition of base from day 40, the ratio increased to 1.33 ± 0.18 g-N g-N^-1^. The effluent also contained nitrate, but represented only 5 ± 1% of the sum of nitrite and nitrate. The pH-based feeding strategy resulted in a HRT and thus a SRT of 7 ± 5 days. Approximately 85% of the soluble COD in the influent was removed regardless of the HRT, resulting in a COD/N ratio of 0.2 g-COD g-N^-1^ in the effluent. For more information on reactor performance, see **SI 5**.

**Fig 3:**
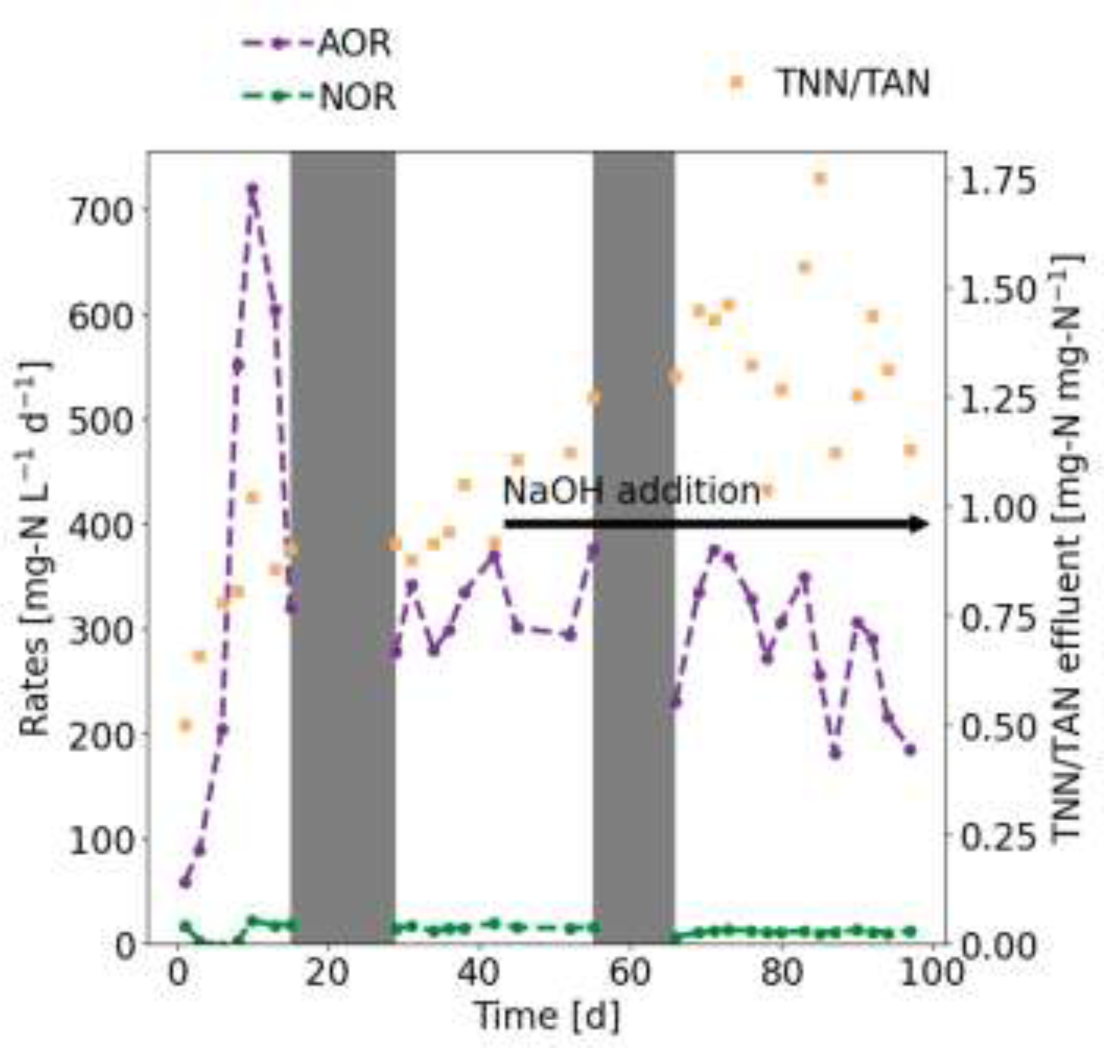
Partial nitritation reactor operated initially without NaOH addition and from day 43 onwards with NaOH addition. AOR = ammonia oxidation rate, NOR = nitrite oxidation rate, TAN = total ammoniacal-nitrogen (NH_3_-N + NH_4_ ^+^-N), TNN = total nitrite-nitrogen (HNO_2_-N + NO_2_ ^-^-N). The grey bars represent two temporary reactor breakdowns most likely due to cleaning agents in the influent.

When the nitritation reactor had been in operation for 55 days, the anoxic reactor was started. Initially, the nitritation rate in the first reactor limited the activity in the anoxic reactor, resulting in nitrogen removal rates of around 300 mg-N L^-1^ d^-1^ (**Fig 4A**). However, around day 80, i.e. after 25 days of operation of the anoxic reactor, the NRR started to decrease to values close to zero by the end of the experiment. The NRE remained high throughout the experiment at 94 ± 1%. The nitrate concentration in the effluent of the anoxic reactor was 129 ± 23 mg-N L^-1^, which was about twice the amount entering the reactor (**Fig 4B**). The nitrite concentration in the effluent was 6 ± 8 mg-N L^-1^, and the ammonium concentration was 26 ± 17 mg-N L^-1^. Salinity increased from 3.5 to 11 mS cm^-1^. The SRT in the anoxic reactor remained above 200 days throughout the experiment. The biomass concentration in the anoxic reactor decreased strongly from initial values around 4 g-VSS L^-1^ to values around 1 g-VSS L^-1^ at the end of the experiment. For more information on reactor performance, see **SI 6**.

**Fig 4:**
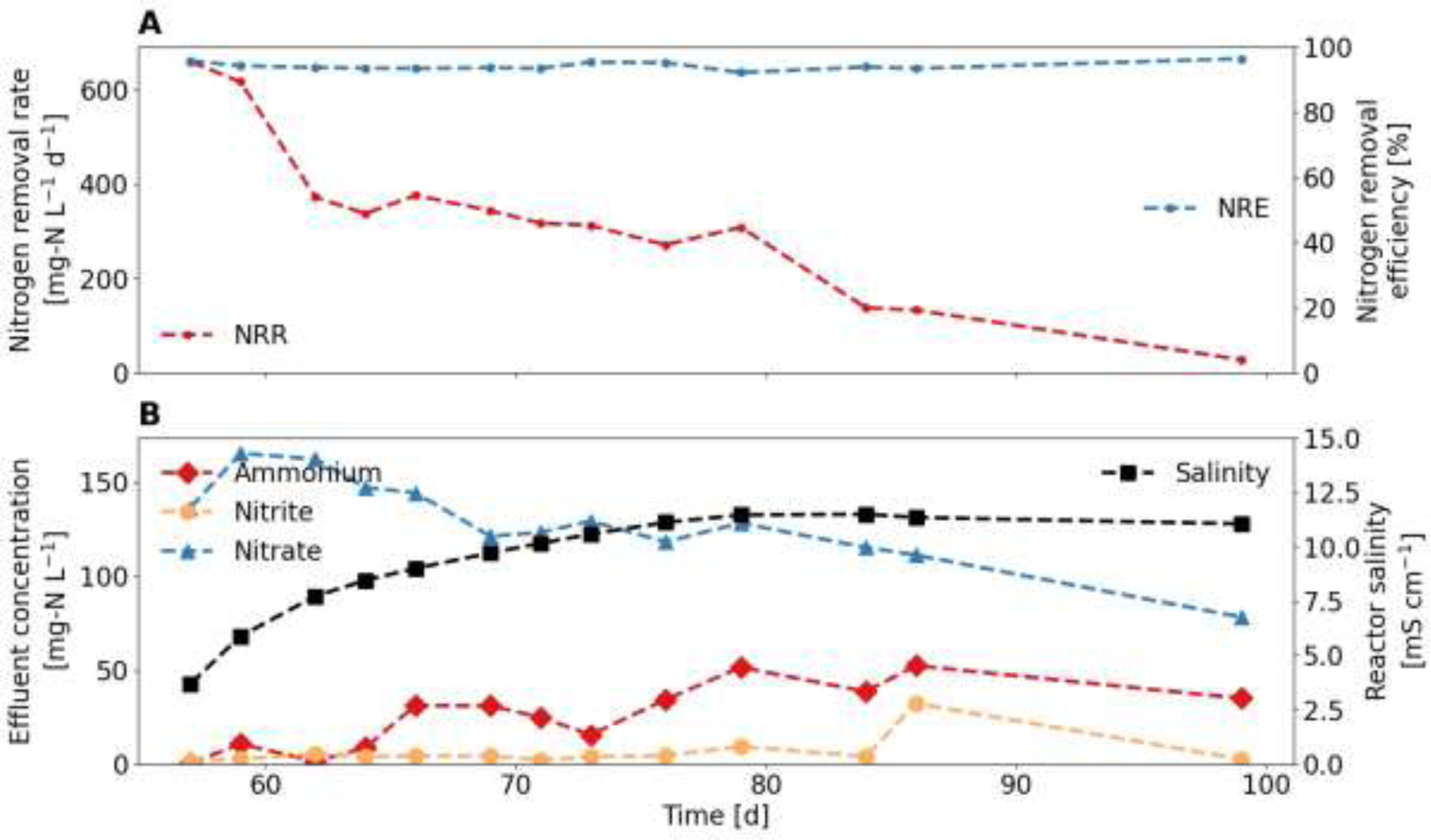
The anoxic reactor was started after the partial nitritation reactor had been running for 55 days. (A) Nitrogen removal rate (NRR) and nitrogen removal efficiency (NRE). (B) Nitrogen compounds and salinity in the effluent.

### 3.2 One-stage partial nitritation/anammox with floccular sludge

#### 3.2.1 Source-separated urine

It was not possible to successfully establish the anammox process using floccular sludge and source-separated urine. After switching the influent from digester supernatant to urine, the NRR initially increased from about 450 mg-N L^-1^ d^-1^ to 600 mg-N L^-1^ d^-1^ in the first week, but then decreased rapidly to about 15 mg-N L^-1^ d^-1^ (**Fig 5A**). After the influent switch, the anoxic feeding phase had to be increased from about 40% to 90% of the feeding phase due to the decreasing AnAOB activity. The calculated SRT increased sharply from 25 days for the digester supernatant to up to 2000 days. Despite the increase in SRT, the biomass concentration remained constant at about 2800 mg-VSS L^-1^. The NRE was between 80 - 90% for both influents. The ammonium concentration in the effluent increased from about 75 mg-N L^-1^ for the digester supernatant to about 200 mg-N L^-1^ for the urine (**Fig 5B**). The peak in ammonium around day 20 was due to a reactor control failure leading to continuous pumping and a pH increase. The nitrite concentration in the effluent remained below 2 mg-N L^-1^ throughout the experiment. The nitrate concentration in the effluent also remained fairly constant at about 100 mg-N L^-1^, and the nitrate production rate decreased along with the NRR. The salinity of the effluent increased from 2 to 10 mS cm^-1^, and the ratio of monovalent to divalent cations increased from 2 mol mol^-1^ to 140 mol mol^-1^. For more information on reactor performance, see **SI 7**.

**Fig 5:**
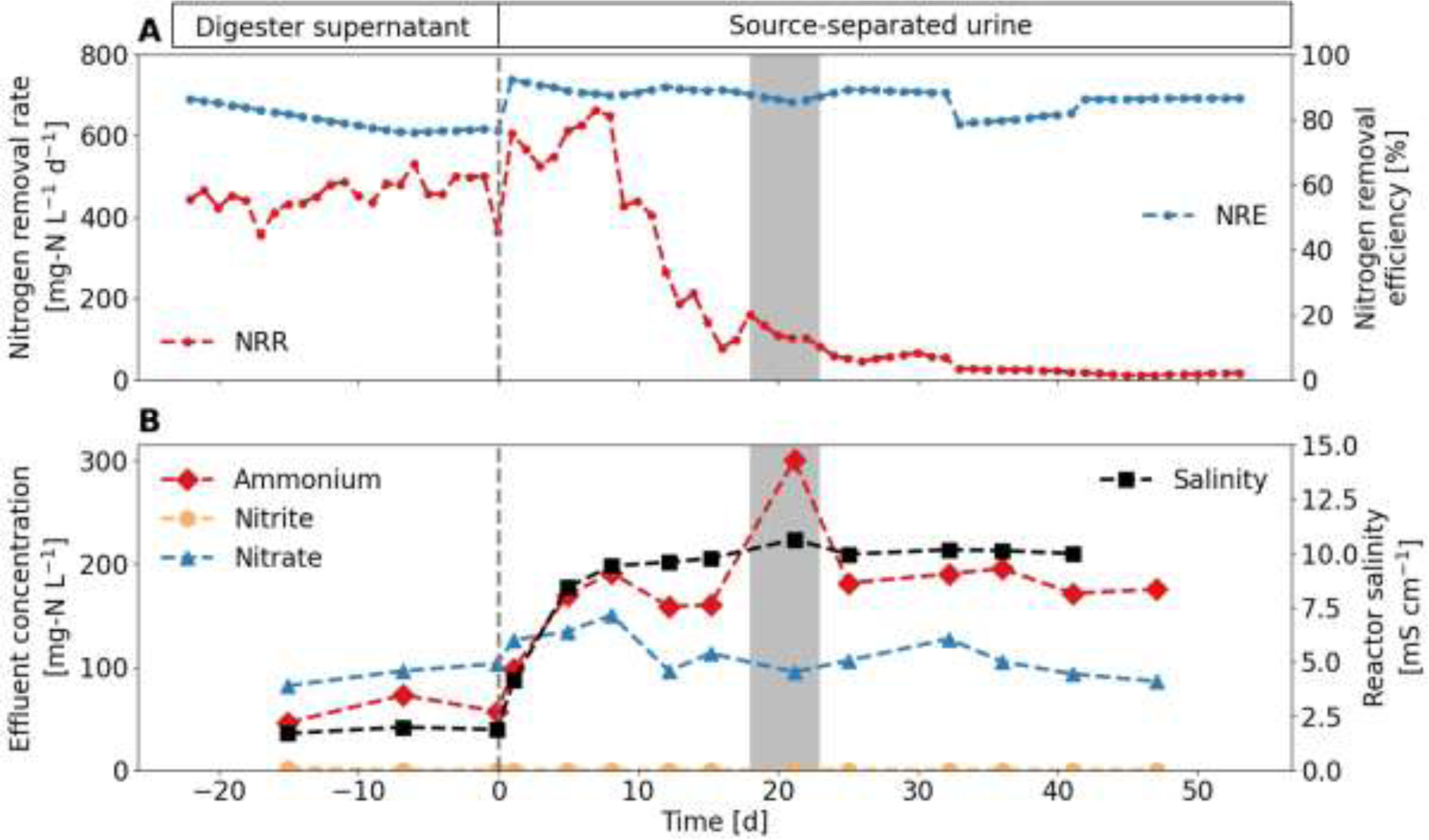
Reactor initially fed with digester supernatant and switched (dotted line) to source-separated urine. (A) Nitrogen removal rate (NRR) and nitrogen removal efficiency (NRE). (B) Nitrogen compounds and salinity in the effluent. A reactor failure from day 18 to day 23 (grey shading) resulted in continuous influent pumping and a pH increase to 8.4 and free ammonia concentrations of 40 mg-N L^-1^.

The dominant AerAOB in the reactor was *Nitrosomonas* sp., NOB consisted of *Nitrospira* sp., *Nitrobacter* sp., and “*Candidatus* Nitrotoga”, and for the AnAOB, “*Candidatus* Brocadia” was found (relative abundance in **SI 7**). The relative abundance of all three functional groups decreased towards the end of the experiment.

#### 3.2.2 Synthetic urine

The PN/A reactor operated with floccular sludge and synthetic urine was not able to maintain the nitrogen removal rate. When synthetic urine was used as an influent, the NRR decreased from about 500 mg-N L^-1^ d^-1^ to 10 mg-N L^-1^ d^-1^ (**Fig 6A**). The calculated SRT increased to about 1000 days and the VSS concentration in the reactor remained constant at about 2300 mg-VSS L^-1^ after the influent switch. After the influent switch, the NRE increased from 80% to 90% in the first week, before dropping to as low as 40% towards the end of the experiment. The ammonium concentration in the effluent increased from about 60 mg-N L^-1^ for the digester supernatant to about 500 mg-N L^-1^ for the urine (**Fig 6B**). The nitrite concentration in the effluent increased from 0 to 15 mg-N L^-1^ after the influent switch. The nitrate concentration in the effluent first decreased after the influent switch and then increased to about 400 mg-N L^-1^. The nitrate production rate strongly increased on day 10 to day 15 (nitrate production rate in **SI 8**). From day 20, the nitrite concentration decreased very slowly during the anoxic phase, and the duration of the feeding phase without aeration was extended. The salinity increased from 2 to 13 mS cm^-1^. For more information on reactor performance, see **SI 8**.

**Fig 6:**
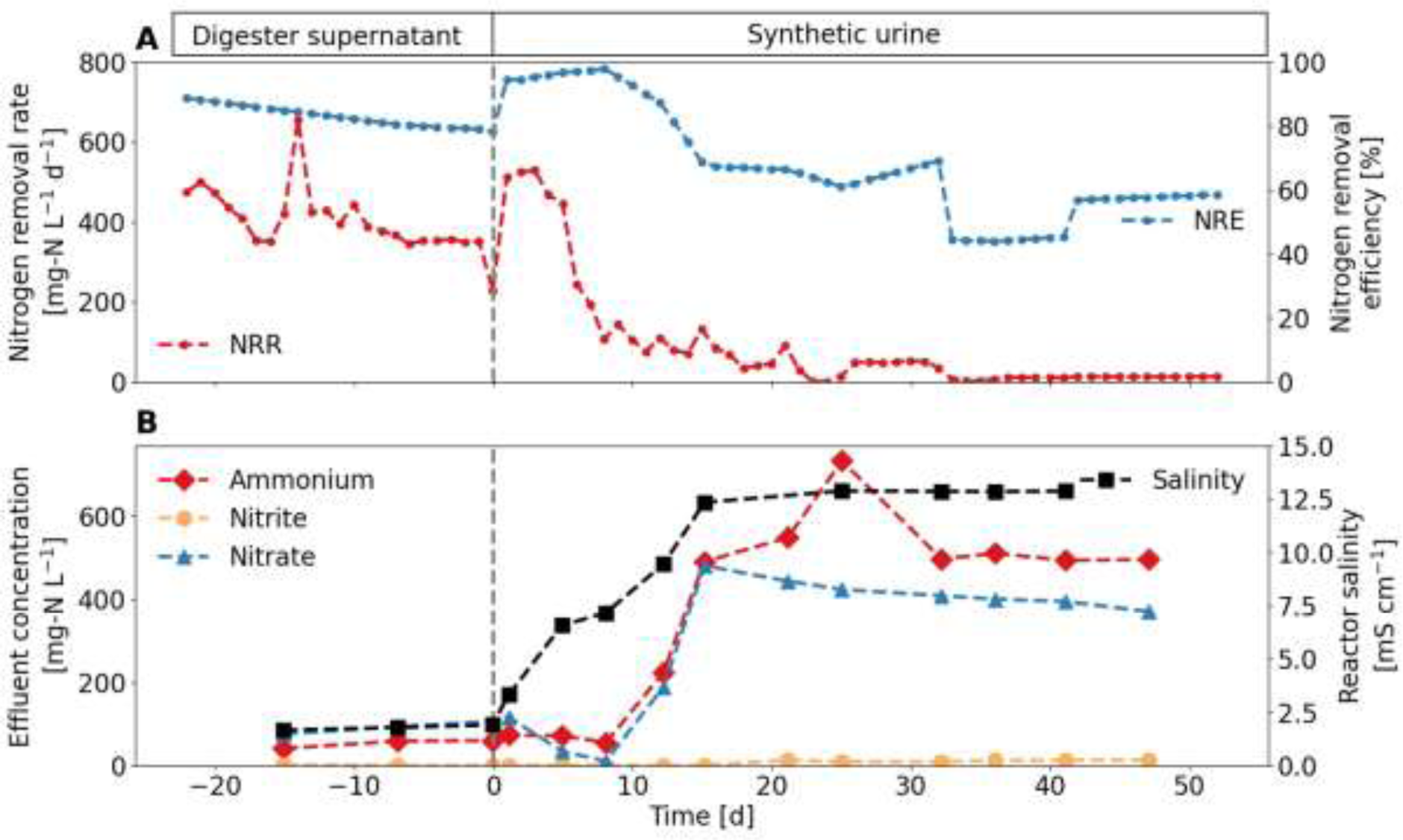
Reactor initially fed with digester supernatant and switched to (dotted line) to synthetic urine. (A) Nitrogen removal rate (NRR) and nitrogen removal efficiency (NRE). (B) Nitrogen compounds in the influent and the reactor.

#### 3.2.3 Source-separated urine with additional divalent cations

The PN/A reactor operated with floccular sludge and source-separated urine with additional divalent cations failed. The influent switch resulted in a sharp decrease in NRR (**Fig 7A**). The calculated SRT increased to values above 100 days. The biomass concentration decreased from 2000 to 1000 mg-VSS L^-1^. The NRE decreased to below 80% towards the end of the experiment despite very long anoxic phases. The ammonium concentration in the effluent increased from about 20 mg-N L^-1^ for the digester supernatant to about 300 mg-N L^-1^ for the urine (**Fig 7B**). Similarly, the nitrate concentration increased from 100 to 350 mg-N L^-1^. On day 10 the nitrate production rate increased despite decreasing NRR (**SI**). The nitrite concentration was mostly below 2 mg-N L^-1^ throughout the experiment. Salinity increased from 2 to 11 mS cm^-1^. The monovalent to divalent cation ratio remained at about 3 mol mol^-1^ after the influent switch due to the addition of divalent cations. During the experiment, the pH set point was gradually reduced from about 7 to 6 to achieve lower concentrations of free ammonia, which did not improve the AnAOB activity. For more information on reactor performance, see **SI 9**.

**Fig 7:**
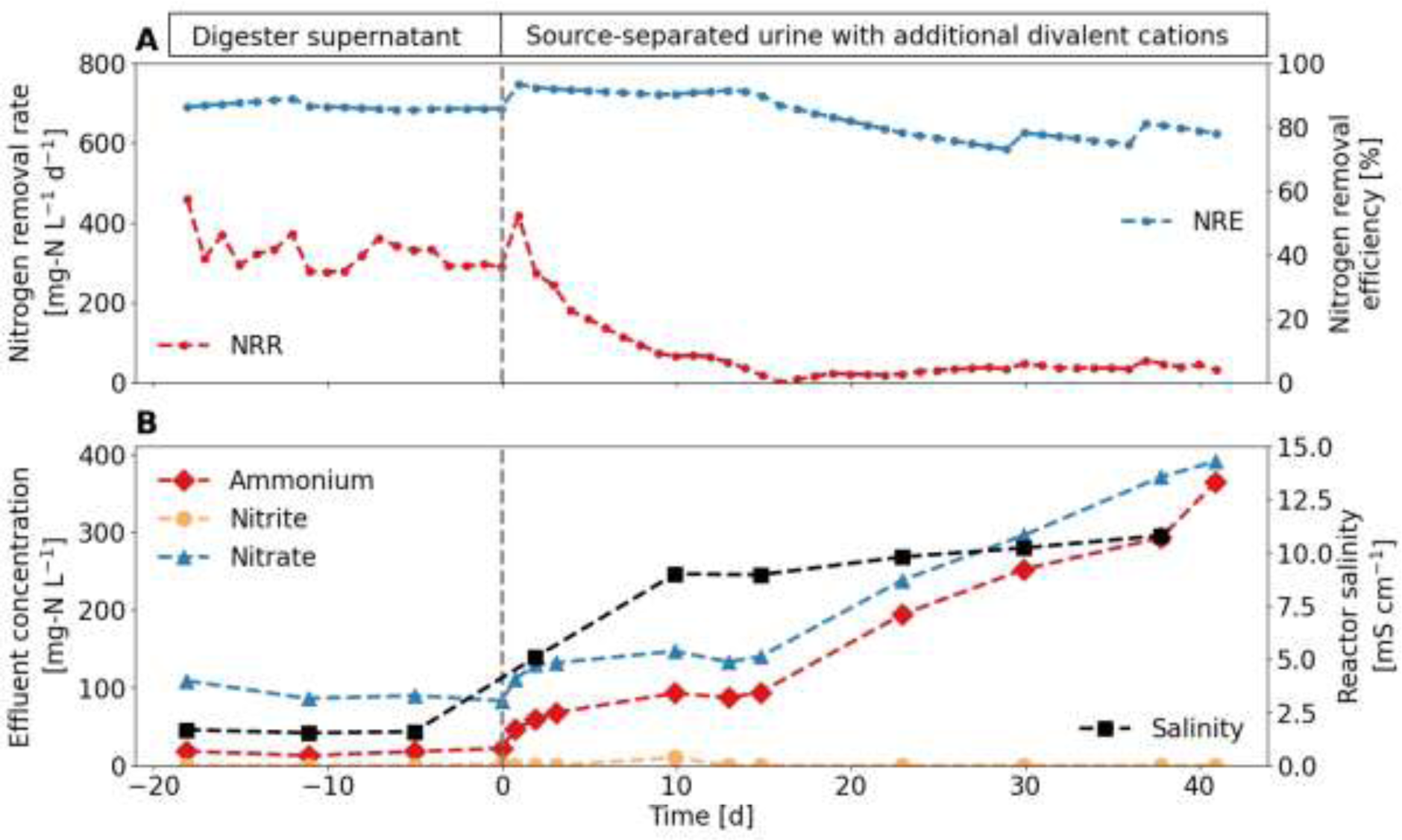
Reactor initially fed with digester supernatant and switched to (dotted line) to source-separated urine with additional divalent cations. (A and B) Nitrogen removal rate (NRR) and nitrogen removal efficiency (NRE). (C and D) Nitrogen compounds in the influent and the reactor.

### 3.3 One-stage partial nitritation/anammox with biofilm carrier

The PN/A reactor operated with biofilm carriers and source-separated urine was successfully operated for 140 days. After the influent switch, the NRR increased to up to 1000 mg-N L^-1^ d^-1^ with an average of 410 ± 220 mg-N L^-1^ d^-1^ (**Fig 8A**). Towards the end of the experiment, the NRR decreased again down to 300 mg-N L^-1^ d^-1^. The NRE was above 90% for most of the time and for both influents, with an average of 93 ± 3% for urine and reaching up to 98% during the last 15 days of the experiments. The short dip to less than 80% around day 28 was caused by a pH control failure which resulted in a continuous influent and TAN concentrations of 450 mg-N L^-1^ and free ammonia concentrations of 40 mg-N L^-1^ in the reactor for several hours, which was subsequently lowered again (**Fig 8B** and **Fig 8C**). Apart from this event, the ammonium concentration in the effluent increased from about 50 mg-N L^-1^ for the digester supernatant to about 150 mg-N L^-1^ for the urine. The nitrite concentration in the effluent was generally less than 2 mg-N L^-1^ throughout the experiment. The nitrate concentration in the effluent was at about 100 mg-N L^-1^ after the influent switch, and the nitrate production correlated with the NRR (nitrate production rate in **SI 10**). The salinity first decreased from 5 mS cm^-1^ in the inoculum to 2 mS cm^-1^ and then increased to 11 mS cm^-1^ after the influent switch. On day 100, the airflow was increased to about to 6.5 L min^-1^ (**Fig 8C**). After the influent switch, the time during which the reactor was not aerated was reduced to as little as 5% of the reaction phase (**Fig 8D**). Towards the end of the experiment, the COD concentration in the influent more than doubled. For more information on reactor performance, see **SI 10**.

**Fig 8:**
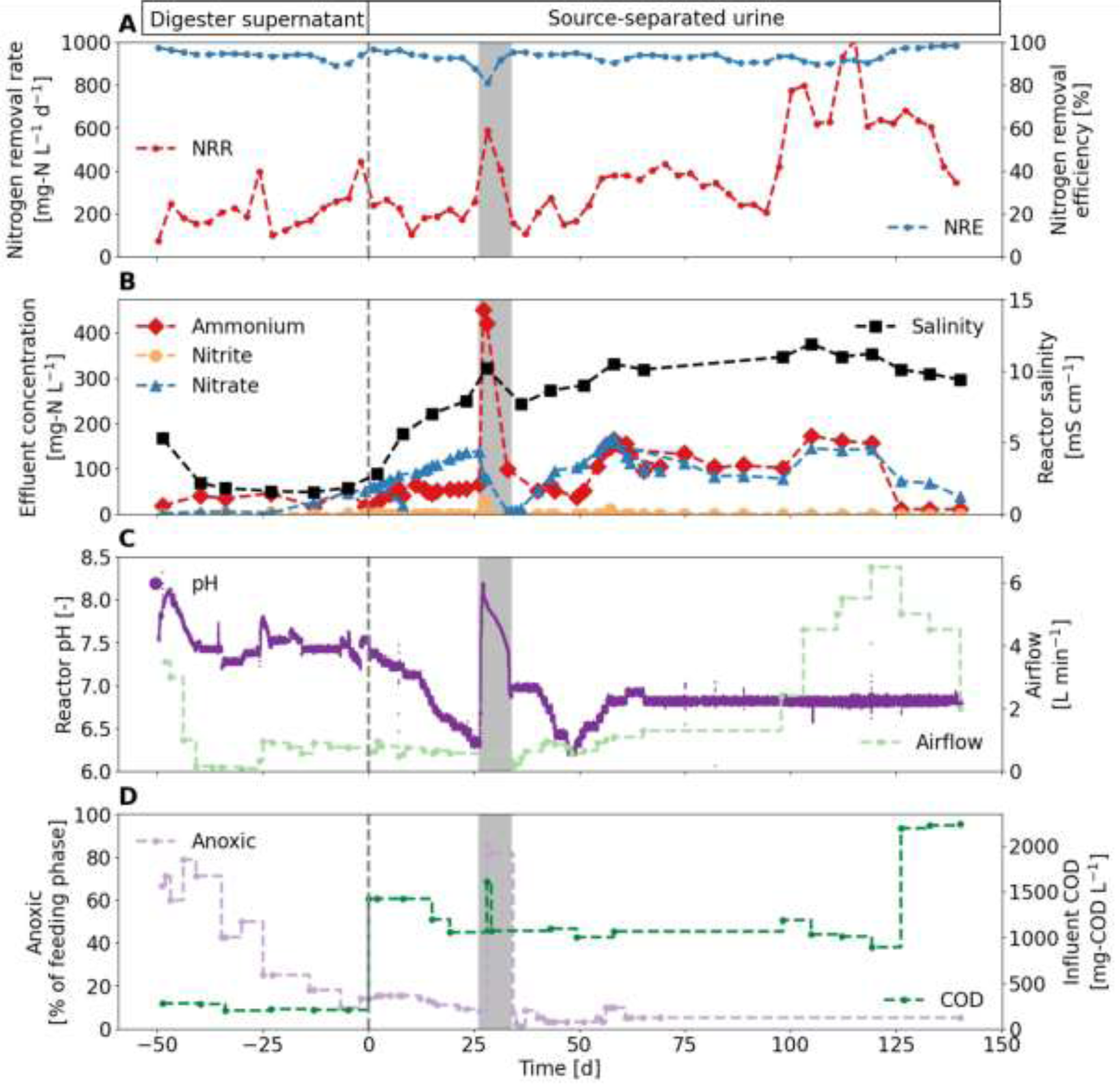
Reactor initially operated with digester supernatant and switched to (dotted line) source-separated urine. (A) Nitrogen removal rate (NRR) and nitrogen removal efficiency (NRE). (B) Nitrogen compounds in the influent and the reactor. (C) pH values. A reactor failure from day 26 to day 34 (grey shading) resulted in continuous influent pumping. (D) Proportion of the feeding phase without aeration and COD/N ratio in the influent.

The dominant AnAOB throughout the experiment was “*Candidatus* Brocadia”, which was mainly found on the biofilm carriers (**Fig 9**). The dominant AerAOB in the reactor was *Nitrosomonas* sp. and was found both, in the floccular sludge and on the biofilm carriers, but with a higher relative abundance in the floccular sludge. NOB consisted of *Nitrospira* sp., *Nitrobacter* sp., and “*Candidatus* Nitrotoga” and low relative abundances were found both, in the floccular sludge and on the biofilm carriers.

**Fig 9:**
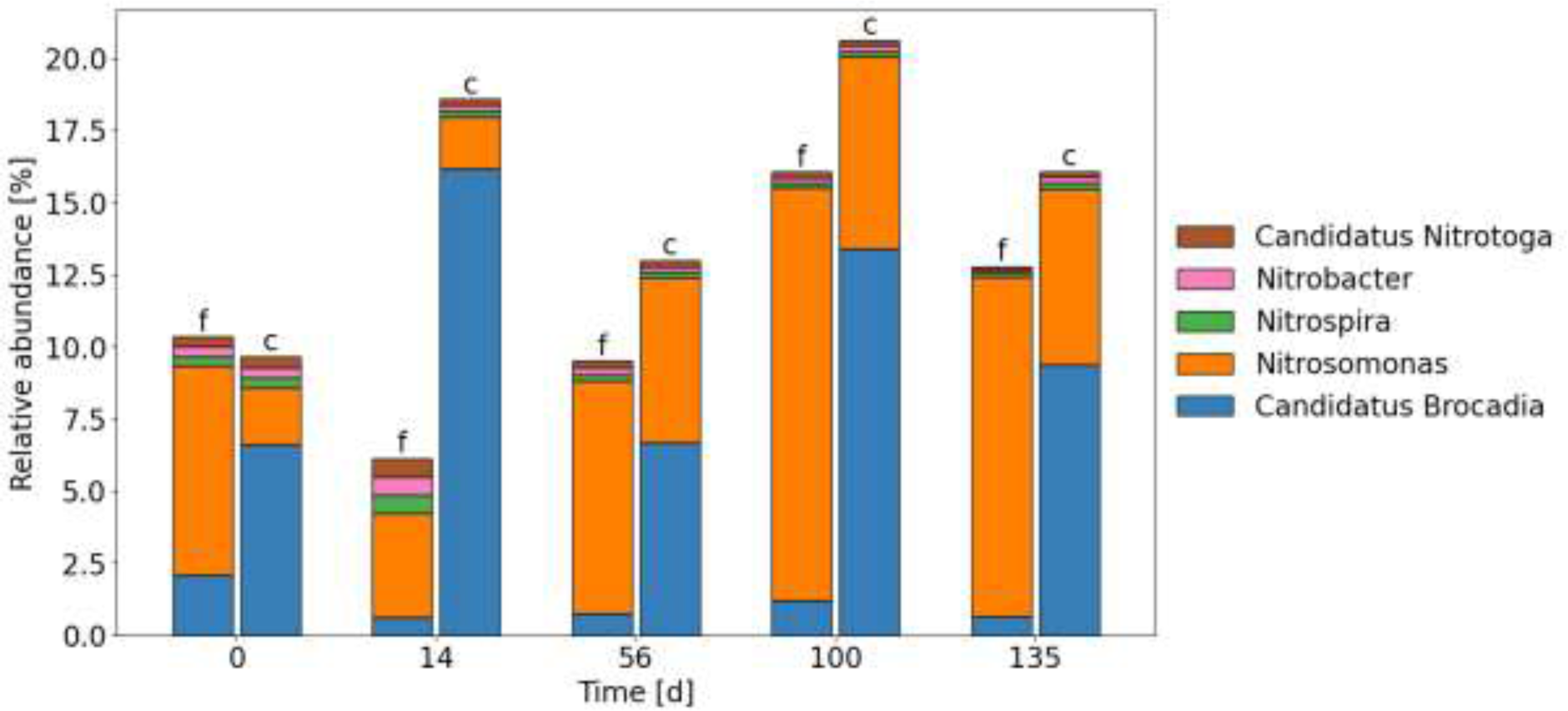
Relative read abundance of recognised genera of AerAOB, NOB, and AnAOB in the floccular sludge (f) and on the biofilm carriers (c).

## 4 Discussion

### 4.1 Successful partial nitritation in two-stage and one-stage configuration using floccular sludge

High ammonia oxidation rates and low nitrite oxidation rates were observed in the two-step and one-step PN/A configurations with the urine influent. The ammonia oxidation rates of 315 ± 130 mg-N L^-1^ d^-1^ in the partial nitritation reactor were in the range observed for urine nitrification and floccular sludge of 430 ± 190 mg-N L^-1^ d^-1^ (Faust et al., 2023b). The low nitrate concentration in the effluent of the partial nitritation reactor indicates only low NOB activity. However, no complete washout was observed at the SRT of about 7 days. According to Faust et al. (2023b), nitrous acid inhibition rather than free ammonia is the main inhibition mechanism for NOB suppression in a partial nitritation reactor operated at a pH of 7. Without the addition of base, the ratio of nitrite to ammonia was 0.9 g-N g-N^-1^, which was below the targeted value of 1.15 to 1.32 g-N g-N^-1^ for anammox (Lotti et al., 2014). Even if NOB were completely inhibited and no nitrite was further oxidized to nitrate, the ratio would only increase to 0.95 mg-N mg-N^-1^. Assuming a theoretical nitrite requirement of 1.32 mole per mole ammonia a maximum of 88% nitrogen could be removed due to the limited alkalinity in urine. The addition of alkalinity allowed more of the ammonia to be converted to nitrate but requires the dosing of a base, i.e., NaOH. Alternatively, an electrochemical unit could be used to produce hydroxide ions on-site (De Paepe et al., 2021).

In the anoxic reactor of the two-stage configuration, the nitrate concentration further increased. NOB activity in the anoxic can be excluded due to the anoxic conditions. Rather, the nitrate concentration increase was a result of anammox matabolism, as about 11% of the converted nitrogen ends up as nitrate as a result of revers electron flow (Strous et al., 1998). Notably, the nitrate increase accounted for only about 5% compared to the converted ammonium and nitrite, meaning that possibly some nitrate was even removed through denitrification. Even though, the organics are mainly degraded in the partial nitritation reactor, some organics could originate from cell debris. Due to the uncertainty of available kinetic data and of the measurements, it is usually not possible to quantify the relative contribution of heterotrophic denitrification and anammox to nitrogen removal (Schielke-Jenni et al., 2015).

In the one-stage reactor fed with source-separated urine, the nitrate in the effluent accounted for less than 11% of the converted nitrogen on average, indicating no or low NOB activity. In addition, the microbial community data showed a decrease in NOB relative abundance during the experiment. For the synthetic and source-separated urine, nitrate production temporarily increased while NRR decreased, indicating potential NOB growth.

### 4.2 Growth of anammox bacteria is the critical step in urine treatment using floccular sludge

While NOB suppression is usually the main challenge in PN/A systems (Hausherr et al., 2022), maintaining AnAOB activity seems to be the main obstacle for urine treatment with floccular sludge. Successful PN/A was possible for digester supernatant in all configurations, but when the influent was changed to source-separated urine, AnAOB activity almost completely ceased within one month. To counteract the decline in AnAOB activity while maintaining high nitrogen removal efficiencies, the anoxic phase was extended, but this did not restore the AnAOB activity. The following section discusses potential factors that may explain the decrease in anammox activity. As AnAOB activity was only lost after the influent switch, the focus of the following discuss will be on the differences between digester supernatant and urine.

#### Toxic organics compounds or pharmaceuticals

Urine contains high concentrations of unidentified organic compounds (Bouatra et al., 2013) and pharmaceuticals (Lienert et al., 2007). However, potentially toxic organic compounds or pharmaceuticals are likely not the reason for the failure of the PN/A process with urine. A strong decrease in AnAOB activity was also observed when the reactor was fed with the synthetic urine containing only readily biodegradable organic substrate and no pharmaceuticals. In the case of the two-stage configuration, the readily biodegradable organics were already degraded before the anammox treatment in the second reactor, but this pretreatment did not prevent failure.

#### COD/N ratio

The availability of biodegradable COD promotes the growth of heterotrophic bacteria, which compete with AnAOB for nitrite and anoxic space. The urine influent had a higher COD/N ratio than the digester supernatant. However, the removal of most of the COD prior to the anoxic reactor in the two-stage configuration did not prevent reactor failure. Furthermore, Jenni et al. (2014) showed that AnAOB are able to coexist with heterotrophic bacteria up to a COD/N ratio in the influent of 1.4 g-COD g-N^-1^, which is higher than in the influent of the one-stage reactors of 0.64 g-COD g-N^-1^. Therefore, the high COD/N ratio can be excluded as the main cause for the failure of AnAOB. However, even if the high COD/N ratio in the urine is not the main cause of failure in the floccular sludge systems, an increase in COD concentration can still have a negative effect on PN/A (see **section 4.3**).

#### Monovalent to divalent cation ratio

The monovalent to divalent cations ratio in urine is with about 350 mol mol^-1^ much higher than in the digester supernatant. Such a high ratio has been reported to negatively affect floc formation and lead to inhibition (Dapena-Mora et al., 2010). However, the addition of divalent cations, i.e. magnesium and calcium, to the urine, which resulted in a monovalent to divalent cation ratio of 13 mol mol^-1^, comparable to that of the digester supernatant of about 15 mol mol^-1^, did not prevent the reactor from failing.

#### Salinity

Urine has a much higher salinity than digester supernatant. Due to the removal of ammonium and bicarbonate during PN/A, the salinity in the reactors is about half of that in the influent. Nevertheless, the electrical conductivity in the reactor increased from about 2 mS cm^-1^ to about 12 mS cm^-1^ after the influent switch. An electrical conductivity of 12 mS cm^-1^ corresponds to 6 g-NaCl L^-1^ and an ionic strength of about 0.12 mol L^-1^ (see **SI 11** for the ionic strength). As shown in **Fig 10A**, there was a negative correlation between the salinity and the nitrogen removal rate for floccular sludge. This negative correlation was not found for the hybrid system, and instead the nitrogen removal rate actually increased (**Fig 10B**).

**Fig 10:**
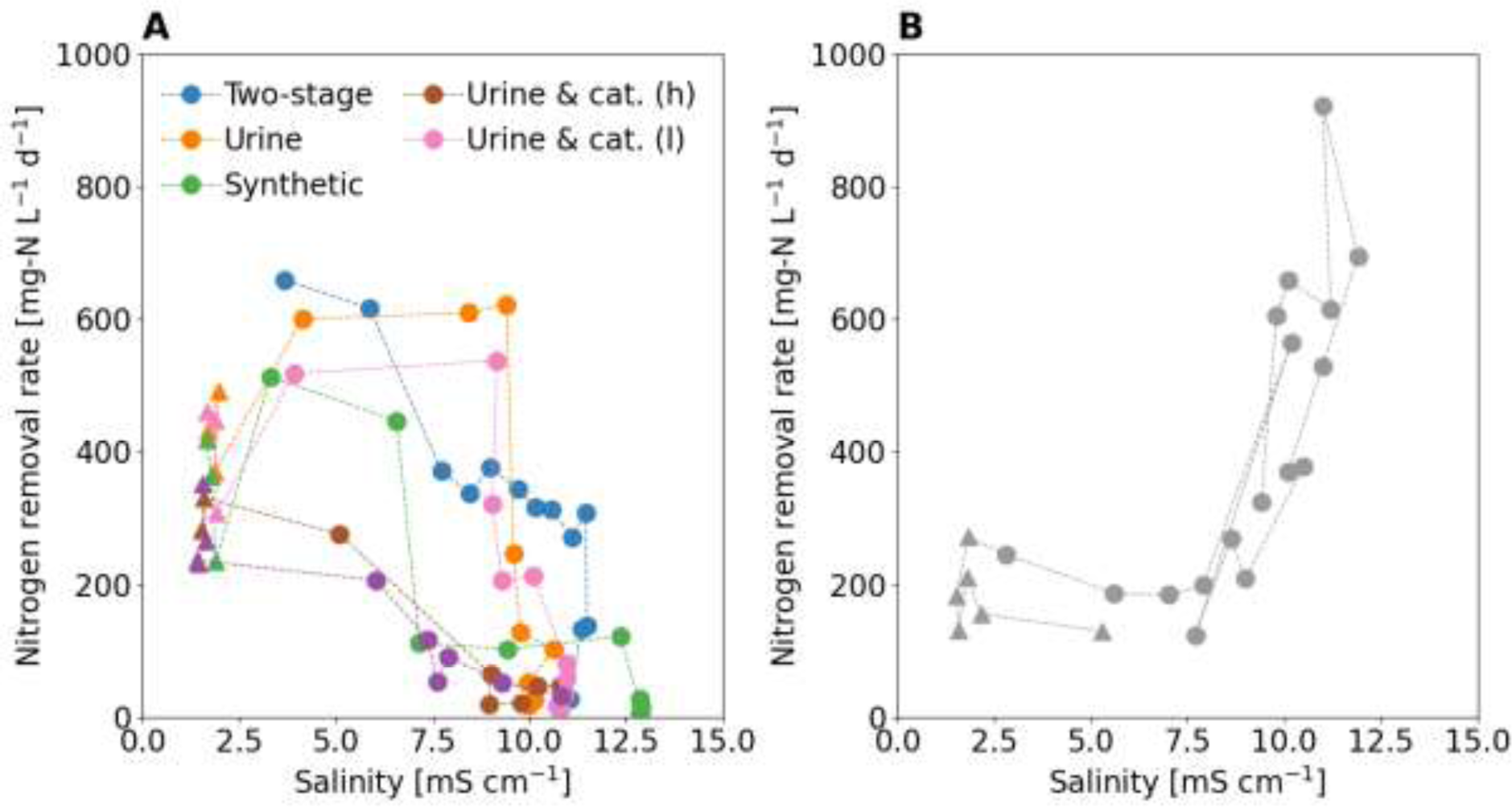
Salinity and nitrogen removal rate. Triangles (▲) represent the datapoints before the influent switch and circles (●) represent the data points after the influent switch. (A) In the reactors operated with floccular sludge, the nitrogen removal rate decreased with increasing salinity. Two additional reactors are shown. One with additional divalent cation but to a lower concentration ("Urine & cat (l)") as in **Section 3.2.3** ("Urine & cat (h)") and one where monovalent cations have been added to the digester supernatant ("Digester & salt") which increased the salinity (see **SI 11** and **SI 12** for additional experiments). (B) In the reactor operated with floccular sludge and biofilm carriers (hybrid), the nitrogen removal rate was maintained or even increased after the influent switch.

Salinity affects the osmotic pressure of the cell membrane which can severely inhibit bacterial metabolism (Hunik et al., 1992). Different inhibition thresholds are reported for different salts, e.g. NaCl or KCl. Dapena-Mora et al. (2010) tested the effect of NaCl on AnAOB activity and reported a half maximal inhibitory concentration (IC_50_) value of 13.5 g-NaCl L^-1^. However, in two research projects, AnAOB activity was maintained at higher NaCl concentrations up to 30 g-NaCl L^-1^ by slowly increasing the salt concentration approximately every 10 days, indicating that AnAOB are able to adapt to high salt concentrations (Kartal et al., 2006; Liu et al., 2009). In the present experiments, the bacteria should have had enough time to adapt to the higher salt concentrations and direct inhibition by salinity seems therefore unlikely.

Salinity also affects floc formation. In case of AnAOB, larger flocs are desirable because they protect AnAOB against oxygen and because they settle better, which improves sludge retention. A favourable affect of salinity on granulation in PN/A systems has been reported (Li et al., 2018). This can be explained with the Derjaguin-Landau-Verwey-Overbeek (DLVO) theory, which states that an increasing salinity reduces electrostatic repulsion by compressing the double layer and therefore improves flocculation (Derjaguin and Landau, 1993). However, according to Zita and Hermansson (1994) this is only true for ionic strengths below 0.1 mol L^-1^. At higher ionic strengths, such as in urine (see **SI 13** for the ionic strength), the high salt content caused the flocs to disintegrate (Zita and Hermansson, 1994). Therefore, a negative effect of salinity on the AnAOB growth in a floccular sludge cannot be excluded.

#### Sulfide

Dapena-Mora et al. (2007) determined an IC_50_ for AnAOB for hydrogen sulfide (H_2_S) of 9.6 mg-S L^-1^. Sulfide can be produced during anaerobic storage of urine by the reduction of sulfate (SO ^2-^), and while the digester supernatant contains low sulfate concentrations below 3 mg L^-1^, the concentration in urine is significant at values of around 500 mg L^-1^ (see **SI 1** for sulfate measurement in influent). However, in the presence of oxygen, sulfide is oxidised very quickly to sulfate. In the case of the two-stage configuration, the sulfide was completely oxidised to sulfate before the solution entered the anoxic reactor. Therefore, an inhibitory effect of sulfide on AnAOB is unlikely.

#### Phosphate

Phosphate has been reported to inhibit AnAOB with an IC_50_ of 650 mg-P L^-1^ (Dapena-Mora et al., 2007). Although the concentration of phosphate in urine was much higher than in the digester supernatant, with concentrations up to 200 mg-P L^-1^, the concentrations were still too low to explain the failure of anammox.

#### Metals

Metals can inhibit AnAOB at high concentrations (Gutwiński et al., 2021), but urine generally contains low concentrations of metals (Fumasoli et al., 2016) making inhibition unlikely. Metals are also essential trace elements for AnAOB growth, and low concentrations lead to limitations (Chen et al., 2014). For example, the growth of acid-tolerant AerAOB in urine was limited by iron (Faust et al., 2022). However, if there were a limiting effect of trace elements, a biofilm system would not solve this problem. Therefore, it is unlikely that metals caused the decrease in activity in the floccular system.

#### Nitrite

Nitrite rather than nitrous acid is the actual inhibiting compound for AnAOB (Lotti et al., 2014). Threshold concentrations are reported to be somewhere between 5 and 280 mg-N L^-1^ (Jin et al., 2012). The nitrite concentration was mostly below 5 mg-N L^-1^ for both, the digester supernatant and the urine and can therefore be excluded.

#### Free ammonia

Free ammonia rather than ammonium was reported to inhibit AnAOB (Dapena-Mora et al., 2007). In short-term experiments, an IC_50_ of 38 mg-N L^-1^ was determined for ammonia and in long-term experiments, ammonia concentration up to 20 mg-N L^-1^ did not significantly affect AnAOB activity (Fernández et al., 2012). With the exception of the reactor failure leading to pH values up to 8.4, the ammonia concentrations in the effluent were always below 10 mg-N L^-1^. Therefore, strong ammonia inhibition on the AnAOB bacteria can be excluded.

#### Sludge retention time

Reported maximum growth rates for AnAOB at around 30°C vary widely (Zhang et al., 2017). Using a rather conservative value of 0.06 d^-1^ (Strous et al., 1998), a minimum SRT of 17 days at optimal conditions is required for the two-stage configuration and of 34 days for the one-stage, considering that the reactor is anoxic 50% of the time. The SRT were much higher in all cases and generally increased after switching from digester supernatant to urine.

#### pH

AnAOB activity was reported to be highest at pH values between 7.0 and 8.5, decreasing to around 40% and 10% at pH values between 6.5 and 6.0. (Strous et al., 1997). However, when operating with highly diluted urine (Bürgmann et al., 2011), a strong increase in AnAOB abundance over time was observed when the pH was decreased to 6.3. In the experiments, neither a constant pH of 7.25 or 7.7 (two-stage) nor a more variable pH between 6 and 7.3 prevented reactor failure. Therefore, pH can likely be excluded as a potential cause of reactor failure.

#### Dissolved oxygen

AnAOB prefer anoxic conditions and can be inhibited by low DO concentrations. However, the DO in the reactor did not change after switching from digester supernatant to urine and remained mostly below 0.2 mg L^-1^. In addition, the DO in the anoxic reactor in the two-stage configuration was essentially zero and yet did not result in successful PN/A. High DO concentrations in the bulk solution can therefore be ruled out as a cause of reactor failure in the presented experiment.

#### Temperature

The optimum range for AnAOB has been reported to be between 30 to 40°C (Jin et al., 2012). The temperature was controlled at a constant temperature of 30°C for both, the digester supernatant and the urine. Therefore, temperature can be excluded as the cause of the decrease in AnAOB activity.

### 4.3 Successful PN/A of urine in one-stage with biofilm carriers

A one-stage PN/A system using floccular sludge and biofilm carriers (hybrid system) was successfully operated for 140 days. The NRR of 410 ± 220 mg-N L^-1^ d^-1^ was in the same range as that reported by Timmer et al. (2023), who achieved an NRR of about of 600 mg-N L^-1^ d^-1^ for stabilised urine with a TN concentration of 2 g-N L^-1^ using a MABR.

Although the influent did not differ from the floccular system, the hybrid system was able to maintain and even increase AnAOB activity. A hybrid system allows for high diffusion resistance of the attached biomass, which protects AnAOB from oxygen and other inhibitory compounds and load fluctuations. For example, the formation of a biofilm can increase microbial resistance to high salinity (Kim and Chong, 2017). A hybrid system also allows for increased sludge retention for the attached biomass. As mentioned in **Section 4.2**, the high salinity of urine can potentially lead to disintegration of the flocs. However, in the hybrid system, the AnAOB grew on the biofilm carriers and were therefore not affected by disintegration, which may explain why the hybrid system was successful. It was even shown that higher salt concentrations increased the production of extracellular polymeric substances on the biofilm, which improved the biofilm structure (Ganesan et al., 2022).

During the experiment with the hybrid system, the anoxic section of the feeding phase was reduced to 5%. The oxygen gradient in the biofilm ensured that AnAOB were not inhibited by oxygen, allowing them to be active even during the aerated feeding phase. An increase in the air flow rate resulted in an increase in the nitrogen removal rate, indicating that nitritation was most likely the rate-limiting step in this configuration. The brief increase in free ammonia up to 40 mg-N L^-1^ on day 18 did not cause any lasting damage to the NRR, even though the concentration is close to the IC_50_ of 38 mg-N L^-1^ (Fernández ^e^t al., 201^2^). This again suggests that the AnAOB may be additionally protected by the biofilm. Finally, an increase in the COD to N ratio in the influent, and therefore the COD load, is likely to have caused a decrease in NRR during the last 15 days. Increased COD is likely to result in the growth of more heterotrophic bacteria competing with AnAOB for space and nitrite, which could lead to a decrease in AnAOB activity.

## 5 Conclusions

- Long-term anammox activity could not be established for source-separated urine in either one-stage or two-stage configurations using floccular sludge.
- Inhibition of AnAOB by toxic organic compounds, whether natural metabolites or pharmaceuticals, has not been confirmed.
- The high salt content, and not the high ratio of monovalent to divalent cations, is probably the main reason for the failure of floccular PN/A with urine.
- It is likely that the high salinity causes large flocs to disintegrate and the AnAOB bacteria to be washed out.
- The use of biofilm carriers (hybrid system) resulted in successful PN/A at high rates and high removal efficiencies.

## Acknowledgments

Philipp Markus, Marijn J. Timmer, and Jolien de Paepe were supported by the European Space Agency (ESA, contract no. 4000130179/20/NL/KML) within the MELiSSA project (www.melissafoundation.com). Sarina Schielke-Jenni was supported by the Swiss National Science (grant No. 200021_125133 and 200020_144498). The authors thank Sylvia Richter (Eawag) and Karin Rottermann (Eawag) for assistance with the chemical analyses and Marco Kipf (Eawag) and Bettina Sterkele (Eawag) for technical support.

## Data availability statement

The raw data of all measured parameters will be made available after acceptance for publication in the following FAIR repository: https://data.eawag.ch/dataset/partial-nitritation-anammox-urine

The original contributions in the form of 16S rRNA data presented can be found under: https://www.ncbi.nlm.nih.gov/sra/PRJNA1051360.

