## Supporting Information for "Conditions for successful nitrogen removal from source-separated urine by partial nitritation/anammox"

#### SUPPLEMENTARY INFORMATION

**Valentin Faust[a,b *], Philipp Markus[c *], Sarina Schielke-Jenni[d], Marijn J. Timmer[e,f], Jolien de Paepe[e,f], Ramon Ganigué[f,g], Siegfried E. Vlaeminck[e,f], Kai M. Udert[a,b **]**

[a] Eawag, Swiss Federal Institute of Aquatic Science and Technology, 8600 Dübendorf, Switzerland

[b] ETH Zürich, Institute of Environmental Engineering, 8093 Zürich, Switzerland

[c] HFS Aqua AG, 8400 Winterthur, Switzerland

[d] AWEL Department for Waste, Water, Energy, and Air, 8090 Zürich, Switzerland

[e] Research Group of Sustainable Energy, Air and Water Technology, Department of Bioscience Engineering, Faculty of Science, University of Antwerp, 2020 Antwerpen, Belgium

[f] Center for Advanced Process Technology for Urban Resource Recovery (CAPTURE), Frieda Saeysstraat 1, 9052 Gent, Belgium

[g] Center for Microbial Ecology and Technology, Department of Biotechnology, Faculty of Bioscience Engineering, Ghent University, Coupure Links 653, 9000 Gent, Belgium

Emails of the authors:

* These authors contributed equally to this work (shared first authorship)

### 1 Influent composition for all experiments

Experiment 1 was started directly with urine. Only TAN, COD and electric conductivity were measured in the influent, which are already reported in **Table 1** of the main manuscript. The reactors in experiments 2 and 3 were both initially fed with the digestion supernatant in **Table S1**.

**Table S1**: Digester supernatant used for the start-up of experiment 2 and 3 (n = 4).

| | | Min | Mean | Max | Stdv. |
|---|---|---|---|---|---|
| Chloride | [mg $L^{-1}$] | 139 | 139 | 140 | 0.5 |
| Phosphate | [mg-P $L^{-1}$] | <1 | <1 | <1 | NA |
| Sulfate | [mg $L^{-1}$] | <3 | <3 | <3 | NA |
| Sodium | [mg $L^{-1}$] | 71 | 74 | 75 | 2 |
| Ammonium | [mg-N $L^{-1}$] | 688 | 697 | 700 | 6 |
| Potassium | [mg $L^{-1}$] | 80 | 82 | 85 | 2 |
| Calcium | [mg $L^{-1}$] | 6 | 78 | 117 | 26 |
| Magnesium | [mg $L^{-1}$] | 42 | 42 | 43 | 0.5 |
| $COD_{sol}$ | [mg-$O_2$ $L^{-1}$] | 171 | 193 | 207 | 15 |
| Conductivity | [mS $cm^{-1}$] | 5.8 | 5.9 | 5.9 | 0.03 |
| pH | [-] | 8.1 | 8.2 | 8.2 | 0.03 |
| Temperature | [°C] | 18 | 19 | 20 | 0.8 |

In experiment 2 the reactor was subsequently fed with source-separated urine shown in **Table S2**.

**Table S2**: Source-separated urine used for experiment 2 (n = 8). *estimated from reactor concentration

| | | Min | Mean | Max | Stdv. |
|---|---|---|---|---|---|
| Chloride | [mg $L^{-1}$] | 2070 | 2350 | 2660 | 180 |
| Phosphate | [mg-P $L^{-1}$] | 118 | 136 | 179 | 20 |
| Sulfate | [mg $L^{-1}$] | 490 | 542 | 626 | 47 |
| Sodium | [mg $L^{-1}$] | 830 | 1170 | 1440 | 170 |
| Ammonium | [mg-N $L^{-1}$] | 1470 | 2410 | 2790 | 470 |
| Potassium | [mg $L^{-1}$] | 770 | 1080 | 1230 | 140 |
| Calcium * | [mg $L^{-1}$] | | 20* | | |
| Magnesium * | [mg $L^{-1}$] | | 10* | | |
| $COD_{sol}$ | [mg-$O_2$ $L^{-1}$] | 770 | 990 | 1820 | 340 |
| Conductivity | [mS $cm^{-1}$] | 19.1 | 21.1 | 24.1 | 1.7 |
| pH | [-] | 8.76 | 8.85 | 8.88 | 0.05 |
| Temperature | [°C] | 18.2 | 20.1 | 21.7 | 1.35 |

For experiment 3 the recipe in **Table S3** was used to produce the synthetic urine in **Table S4**.

**Table S3**: Synthetic urine formula. The synthetic urine formula was designed to mimic stored urine after urea hydrolysis, but the organic matter was replaced by acetate. In addition, the very low calcium and magnesium concentrations were neglected. An Excel spreadsheet and PHREEQC software were used to determine the formula. The exact formula for the first 15 litres of synthetic urine is shown here as an example. Subsequent batches of synthetic urine were adjusted to the ammonium concentration of the source-separated urine.

| | **Formula** | **Unit** | **Value** |
|---|---|---|---|
| Ammonia-hydrogen-carbonate | $NH_4HCO_3$ | [g $L^{-1}$] | 8.98 |
| Ammonia-hydroxide 32% | $NH_3$ | [ml $L^{-1}$] | 5.95 |
| Ammonia-acetate | $NH_4CH_3COO$ | [g $L^{-1}$] | 2.17 |
| Potassium-chloride | KCl | [g $L^{-1}$] | 2.21 |
| Sodium-di-hydrogen-phosphate-dihydrate | $NaH_2PO_4.2H_2O$ | [g $L^{-1}$] | 0.917 |
| Sodium-chloride | NaCl | [g $L^{-1}$] | 2.69 |
| Sodium-sulfate-decahydrate | $Na2SO_4.10H_2O$ | [g $L^{-1}$] | 1.95 |

**Table S4**: Composition of the synthetic urine used for experiment 3 (n = 6). No magnesium or calcium was added to the synthetic urine.

| | | Min | Mean | Max | Stdv. |
|---|---|---|---|---|---|
| Chloride | [mg $L^{-1}$] | 2650 | 2750 | 2840 | 80 |
| Phosphate | [mg-P $L^{-1}$] | 178 | 185 | 196 | 7 |
| Sulfate | [mg $L^{-1}$] | 568 | 587 | 605 | 15 |
| Sodium | [mg $L^{-1}$] | 980 | 1340 | 1510 | 180 |
| Ammonium | [mg-N $L^{-1}$] | 1480 | 2440 | 2990 | 610 |
| Potassium | [mg $L^{-1}$] | 770 | 1060 | 1180 | 150 |
| Calcium | [mg $L^{-1}$] | 0 | 0 | 0 | 0 |
| Magnesium | [mg $L^{-1}$] | 0 | 0 | 0 | 0 |
| $COD_{sol}$ | [mg-$O_2$ $L^{-1}$] | 1220 | 1570 | 1910 | 340 |
| Conductivity | [mS $cm^{-1}$] | 18.5 | 20.8 | 22.3 | 2.1 |
| pH | [-] | 9.1 | 9.1 | 9.1 | 0.02 |
| Temperature | [°C] | 19 | 20 | 20 | 1 |

The reactor in experiment 4 was initially fed with the digestion supernatant in **Table S5**.

**Table S5**: Digester supernatant used for the start-up of experiment 4 (n = 3).

| | | Min | Mean | Max | Stdv. |
|---|---|---|---|---|---|
| Chloride | [mg $L^{-1}$] | 146 | 152 | 160 | 7 |
| Phosphate | [mg-P $L^{-1}$] | <1 | <1 | <1 | NA |
| Sulfate | [mg $L^{-1}$] | <3 | <3 | <3 | NA |
| Sodium | [mg $L^{-1}$] | 81 | 90 | 102 | 11 |
| Ammonium | [mg-N $L^{-1}$] | 660 | 722 | 830 | 94 |
| Potassium | [mg $L^{-1}$] | 100 | 106 | 116 | 9 |
| Calcium | [mg $L^{-1}$] | 39 | 55 | 75 | 18 |
| Magnesium | [mg $L^{-1}$] | 39 | 44 | 50 | 6 |
| $COD_{sol}$ | [mg-$O_2$ $L^{-1}$] | 205 | 216 | 237 | 18 |
| Conductivity | [mS $cm^{-1}$] | 5.6 | 5.8 | 6.0 | 0.2 |
| pH | [-] | 8.13 | 8.21 | 8.31 | 0.09 |
| Temperature | [°C] | 21.0 | 21.8 | 22.3 | 0.7 |

In experiment 4 the reactor was subsequently fed with source-separated urine with additional divalent cations shown in **Table S6**.

**Table S6**: Source-separated urine with additional divalent cations used for experiment 4 (n = 8). *estimated from reactor concentration

| | | Min | Mean | Max | Stdv. |
|---|---|---|---|---|---|
| Chloride | [mg $L^{-1}$] | 1900 | 2100 | 2380 | 19 |
| Phosphate | [mg-P $L^{-1}$] | 98 | 119 | 144 | 19 |
| Sulfate | [mg $L^{-1}$] | 300 | 485 | 633 | 117 |
| Sodium | [mg $L^{-1}$] | 740 | 1030 | 1170 | 150 |
| Ammonium | [mg-N $L^{-1}$] | 1780 | 2450 | 3090 | 480 |
| Potassium | [mg $L^{-1}$] | 740 | 1030 | 1150 | 150 |
| Calcium* | [mg $L^{-1}$] | | 250 | | |
| Magnesium* | [mg $L^{-1}$] | | 200 | | |
| $COD_{sol}$ | [mg-$O_2$ $L^{-1}$] | 1000 | 1140 | 1610 | 210 |
| Conductivity | [mS $cm^{-1}$] | 18.2 | 21.0 | 24.7 | 2.7 |
| pH | [-] | 8.64 | 8.85 | 8.96 | 0.12 |
| Temperature | [°C] | 22.3 | 24.9 | 28.5 | 1.8 |

The reactor in experiment 5 was initially fed with the digestion supernatant in **Table S7**.

**Table S7**: Digester supernatant used for the start-up of experiment 5 with additional biocarriers (n = 6).

| | | Min | Mean | Max | Stdv. |
|---|---|---|---|---|---|
| Chloride | [mg $L^{-1}$] | 146 | 163 | 181 | 11 |
| Phosphate | [mg-P $L^{-1}$] | <1 | <1 | <1 | NA |
| Sulfate | [mg $L^{-1}$] | <3 | <3 | <3 | NA |
| Sodium | [mg $L^{-1}$] | 81 | 92 | 102 | 8 |
| Ammonium | [mg-N $L^{-1}$] | 678 | 722 | 830 | 58 |
| Potassium | [mg $L^{-1}$] | 99 | 108 | 116 | 7 |
| Calcium | [mg $L^{-1}$] | 42 | 67 | 81 | 16 |
| Magnesium | [mg $L^{-1}$] | 38 | 44 | 50 | 4 |
| $COD_{sol}$ | [mg-$O_2$ $L^{-1}$] | 201 | 230 | 278 | 35 |
| Conductivity | [mS $cm^{-1}$] | 5.7 | 6.0 | 6.1 | 0.2 |
| pH | [-] | 7.9 | 8.1 | 8.3 | 0.1 |
| Temperature | [°C] | 22.6 | 22.0 | 23.2 | 1.1 |

In experiment 5 the reactor was subsequently fed with source-separated urine shown in **Table S8**.

**Table S8**: Source-separated urine used for experiment 5 with additional biocarriers (n = 19). *estimated from reactor concentration

| | | Min | Mean | Max | Stdv. |
|---|---|---|---|---|---|
| Chloride | [mg $L^{-1}$] | 1800 | 2180 | 2860 | 270 |
| Phosphate | [mg-P $L^{-1}$] | 95 | 123 | 160 | 20 |
| Sulfate | [mg $L^{-1}$] | 300 | 490 | 690 | 100 |
| Sodium | [mg $L^{-1}$] | 740 | 1130 | 1430 | 170 |
| Ammonium | [mg-N $L^{-1}$] | 1760 | 2510 | 3230 | 470 |
| Potassium | [mg $L^{-1}$] | 740 | 1120 | 1430 | 170 |
| Calcium* | [mg $L^{-1}$] | | 15 | | |
| Magnesium* | [mg $L^{-1}$] | | 8 | | |
| $COD_{sol}$ | [mg-$O_2$ $L^{-1}$] | 890 | 1360 | 2240 | 450 |
| Conductivity | [mS $cm^{-1}$] | 8.2 | 21.0 | 24.7 | 3.9 |
| pH | [-] | 8.6 | 8.8 | 9.0 | 0.1 |
| Temperature | [°C] | 21.7 | 24.4 | 28.5 | 1.8 |

#### 2 13-L reactor set-up of one-stage reactors

The 13-L reactors are shown in **Figure S1**. The reactors were fed from the bottom with a peristaltic pump (Ecoline, Ismatec®; Masterflex L/S, model 7518-10, Easy-load®). For the effluent, the reactors had an overflow at the top. The reactors were equipped with an overhead stirrer (RZR 1 Overhead Stirrer, Heidolph), a pressure gauge (Cerabar T PMC131, Endress+Hauser), a pH sensor (Orbisint CPS11D, Endress+Hauser), and a DO sensor (Oxymax COS61D, Endress+Hauser). The temperature was also measured with the pH sensor (Orbisint CPS11D, Endress+Hauser) and controlled by a water heat jacket (F32, Julabo).

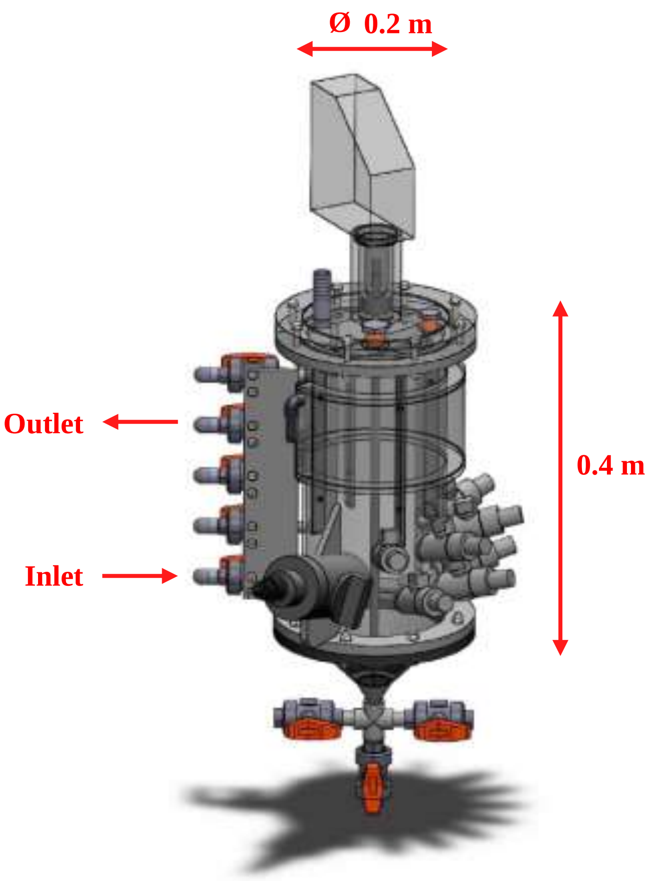

**Figure S1:** Dimension of the 12-L lab scale reactors (picture © Adriano Joss, Eawag).

### 3 Acid-base equilibrium for ammonium and ammonia

The acid-base equilibrium for ammonium and ammonia is shown in **Equation S1**.

$$NH_4^+ \rightleftharpoons NH_3 + H^+ \qquad (S1)$$

Equilibrium concentrations were calculated using **Equations S2**, **S3** (Crittenden et al., 2012), **S4** (Anthonisen et al., 1976), **S5** (Lewis and Randall, 1921). and **S6** (Davies, 1967) following the description in Crittenden et al. (2012).

$$[TAN] = [NH_3] + [NH_4^+] \qquad (S2)$$

$$K_a(T) = \frac{[NH_3] \times [H^+] \times f_{mono}}{[NH_4^+] \times f_{mono}} = \frac{[NH_3] \times 10^{-pH}}{[NH_4^+] \times f_{mono}} \qquad (S3)$$

$$K_a(T) = e^{\frac{-6344}{T}} \qquad (S4)$$

$$I = \frac{1}{2} \times \sum_i [C_i] \times Z_i^2 \qquad (S5)$$

$$\log_{10} f_{mono} = -A * \left( \frac{I^{0.5}}{1 + I^{0.5}} - 0.3 * I \right) \qquad (S6)$$

All variables are shown in **Table S9**.

**Table S9:** All variables used in **Equations S1** to **S6**.

| **Variable** | **Name** | **Unit** |
|---|---|---|
| [TAN] | Total ammoniacal nitrogen concentration | [mol $L^{-1}$] |
| [$NH_3$] | Ammonia concentration | [mol $L^{-1}$] |
| [$NH_4^+$] | Ammonium concentration | [mol $L^{-1}$] |
| [$H^+$] | Proton concentration | [mol $L^{-1}$] |
| $K_a$ | Dissociation constant, $pK_a$=9.1 at 30°C | [mol $L^{-1}$] |
| $f_{mono}$ | Activity coefficient for monovalent ions | [-] |
| T | Absolute temperature, 303 K at 30°C | [K] |
| I | Ionic strength | [mol $L^{-1}$] |
| [$C_i$] | Concentration of ionic specie i | [mol $L^{-1}$] |
| $Z_i$ | Charge of ionic specie i | [-] |
| A | 0.51 at 30°C (Stumm and Morgan, 1996) | [-] |

For the calculation of the ionic strength, the main ionic species in nitrified urine were considered: $K^+$, $Cl^-$, $Na^+$, $NH_4^+$, $NO_3^-$, $NO_2^-$, $H_2PO_4^-$, and $SO_4^{2-}$. The ionic strengths calculated for this publication ranged from 0.05 M to 0.2 M. For ionic strengths below 0.1 M, the Davies equations (**Equation S6**) typically have an error of 1.5%, and for ionic strengths between 0.1 M and 0.5 M an error of 5% to 10% can be expected (Levine, 1988).

### 4 Acid-base equilibrium for nitrous acid and nitrite

The acid-base equilibrium for nitrous acid and ammonia is shown in **Equation S7**.

$$HNO_2 \leftrightarrow NO_2^- + H^+ \qquad (S7)$$

Equilibrium concentrations were calculated using **Equations S8**, **S9** (Crittenden et al., 2012), **S10** (Anthonisen et al., 1976), **S11** (Lewis and Randall, 1921), and **S12** (Davies, 1967), following the description in Crittenden et al. (2012).

$$[TNN] = [HNO_2] + [NO_2^-] \qquad (S8)$$

$$K_a(T) = \frac{[NO_2^-] \times f_{mono} \times [H^+] \times f_{mono}}{[HNO_2]} = \frac{[NO_2^-] \times f_{mono} \times 10^{-pH}}{[HNO_2]} \qquad (S9)$$

$$K_a(T) = e^{\frac{-2300}{T}} \qquad (S10)$$

$$I = \frac{1}{2} \times \sum_i [C_i] \times Z_i^2 \qquad (S11)$$

$$\log_{10} f_{mono} = -A * \left(\frac{I^{0.5}}{1 + I^{0.5}} - 0.3 * I\right) \qquad (S12)$$

All variables are shown in **Table S10**.

**Table S10:** All variables used in **Equations S7** to **S12**.

| Variable | Name | Unit |
|---|---|---|
| [TNN] | Total nitrite nitrogen concentration | [mol $L^{-1}$] |
| [$HNO_2$] | Nitrous acid concentration | [mol $L^{-1}$] |
| [$NO_2^-$] | Nitrite concentration | [mol $L^{-1}$] |
| [$H^+$] | Proton concentration | [mol $L^{-1}$] |
| $K_a$ | Dissociation constant, $pK_a$=3.3 at 30°C | [mol $L^{-1}$] |
| $f_{mono}$ | Activity coefficient for monovalent ions | [-] |
| T | Absolute temperature, 303 K at 30°C | [K] |
| I | Ionic strength | [mol $L^{-1}$] |
| [$C_i$] | Concentration of ionic specie i | [mol $L^{-1}$] |
| $Z_i$ | Charge of ionic specie i | [-] |
| A | 0.51 at 30°C (Stumm and Morgan, 1996) | [-] |

For the calculation of the ionic strength, the main ionic species in nitrified urine were considered: $K^+$, $Cl^-$, $Na^+$, $NH_4^+$, $NO_3^-$, $NO_2^-$, $H_2PO_4^-$, and $SO_4^{2-}$. The ionic strengths calculated for this publication ranged from 0.05 M to 0.2 M. For ionic strengths below 0.1 M, the Davies equations (**Equation S12**) typically have an error of 1.5%, and for ionic strengths between 0.1 M and 0.5 M an error of 5% to 10% can be expected (Levine, 1988)

### 5 Reactor performance of two-stage PN/A: partial nitritation

The nitrogen compounds in the effluent of the nitritation reactor from the two-stage configuration are shown in **Figure S2**. The COD removal and the ratio of organic COD to nitrogen (N) is shown in **Figure S3**.

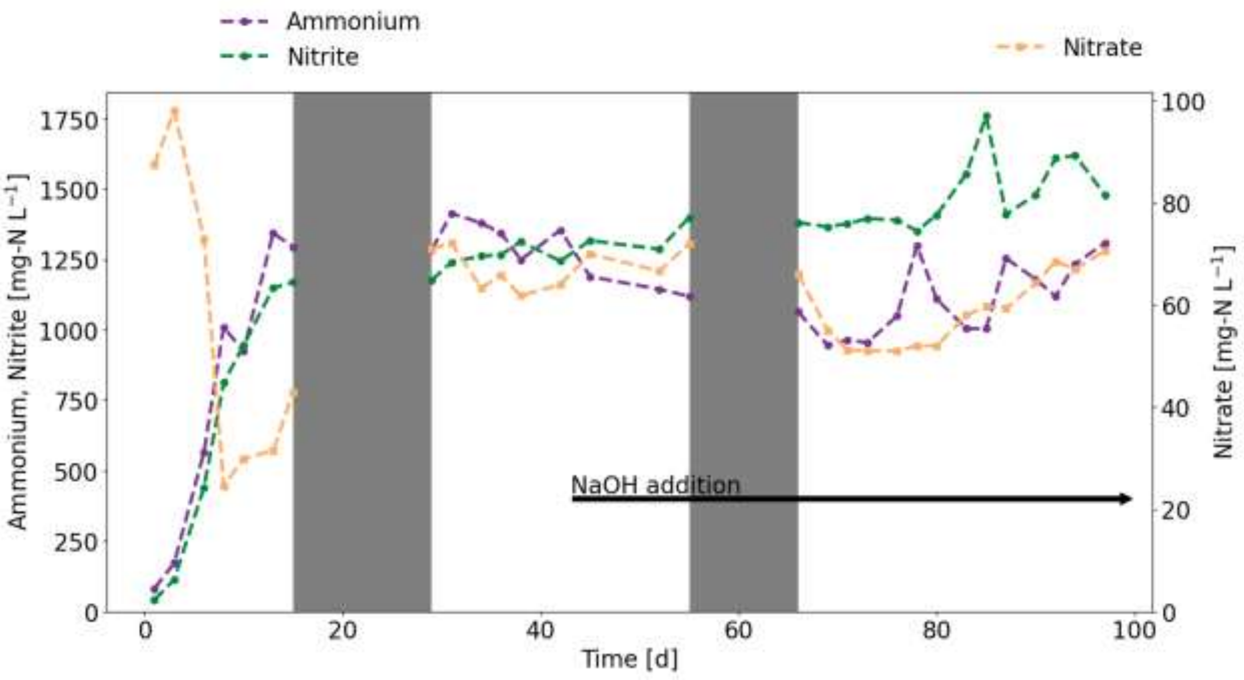

**Figure S2**: Effluent concentration of ammonium, nitrite, and nitrate from the nitritation reactor. The grey bars represent two temporary reactor breakdowns most likely due to cleaning agents in the influent.

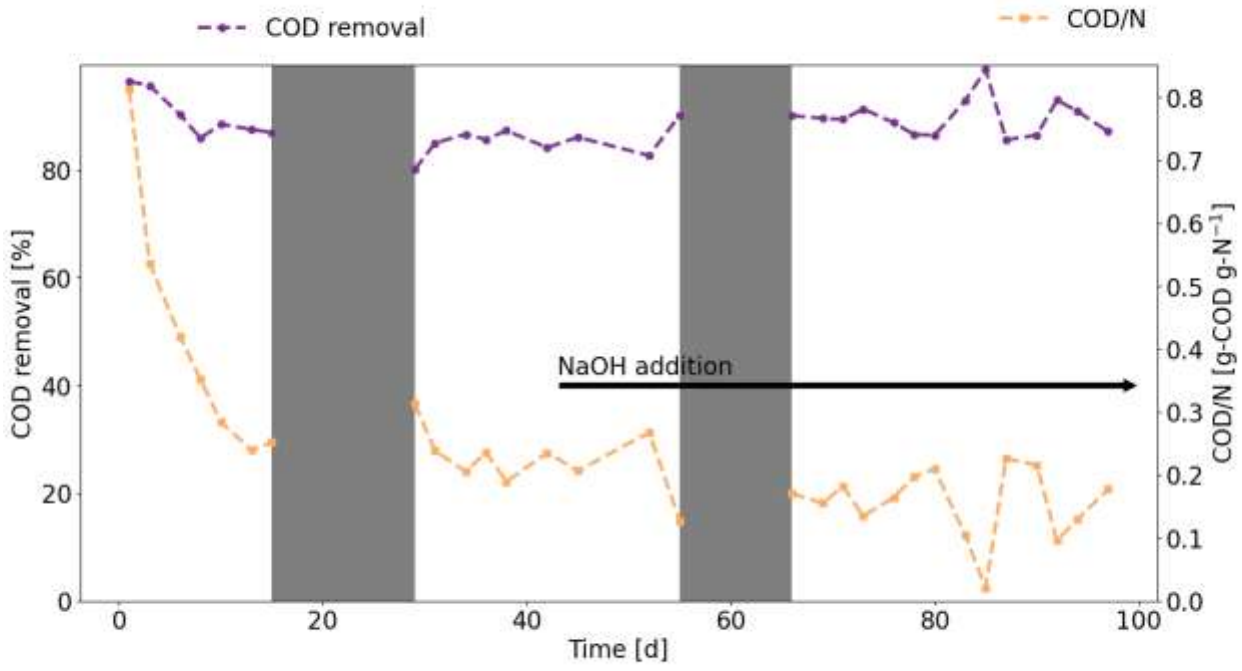

**Figure S3**: COD removal in the nitritation reactor and COD/N ratio in the effluent. The grey bars represent two temporary reactor breakdowns most likely due to cleaning agents in the influent.

### 6 Reactor performance of two-stage PN/A: anammox

The volatile suspended solids (VSS) and the solid retention time (SRT) of the anammox reactor are shown in **Figure S4**.

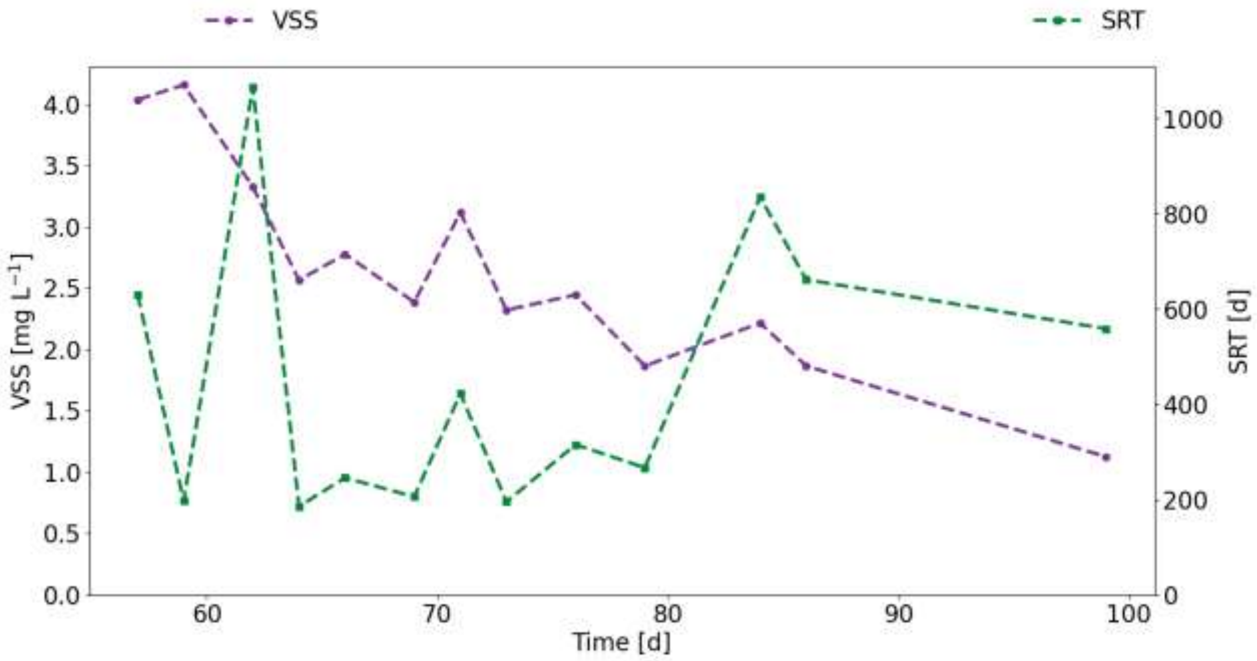

**Figure S4**: Volatile suspended solids (VSS) and the solid retention time (SRT) of the anammox reactor.

### 7 Reactor performance of one stage PN/A: source-separated urine

**Figure S5** shows how much of the feeding phase was performed without aeration. **Figure S6A** shows the VSS concentration in the reactor, **Figure S6B** the solid retention time (SRT), **Figure S6C** the COD degradation comparing the influent and reactor COD, **Figure S6D** the nitrate production rate, **Figure S6E** the monovalent to divalent cations ratio in the reactor, **Figure S6F** the sulfate and phosphate concentrations in the reactor, **Figure S6G** the pH in the reactor, **Figure S6H** the dissolved oxygen (DO) concentration in the reactor.

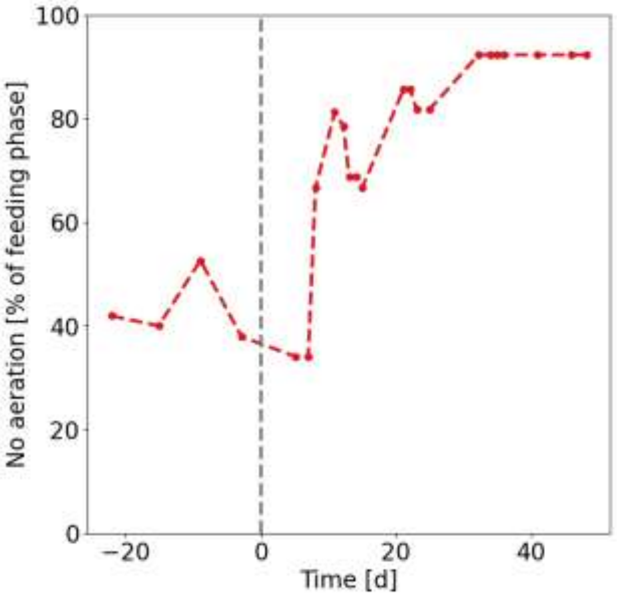

**Figure S5**: After switching from digester supernatant to source-separated urine, the unaerated conditions had to be extended due to declining AnAOB activity.

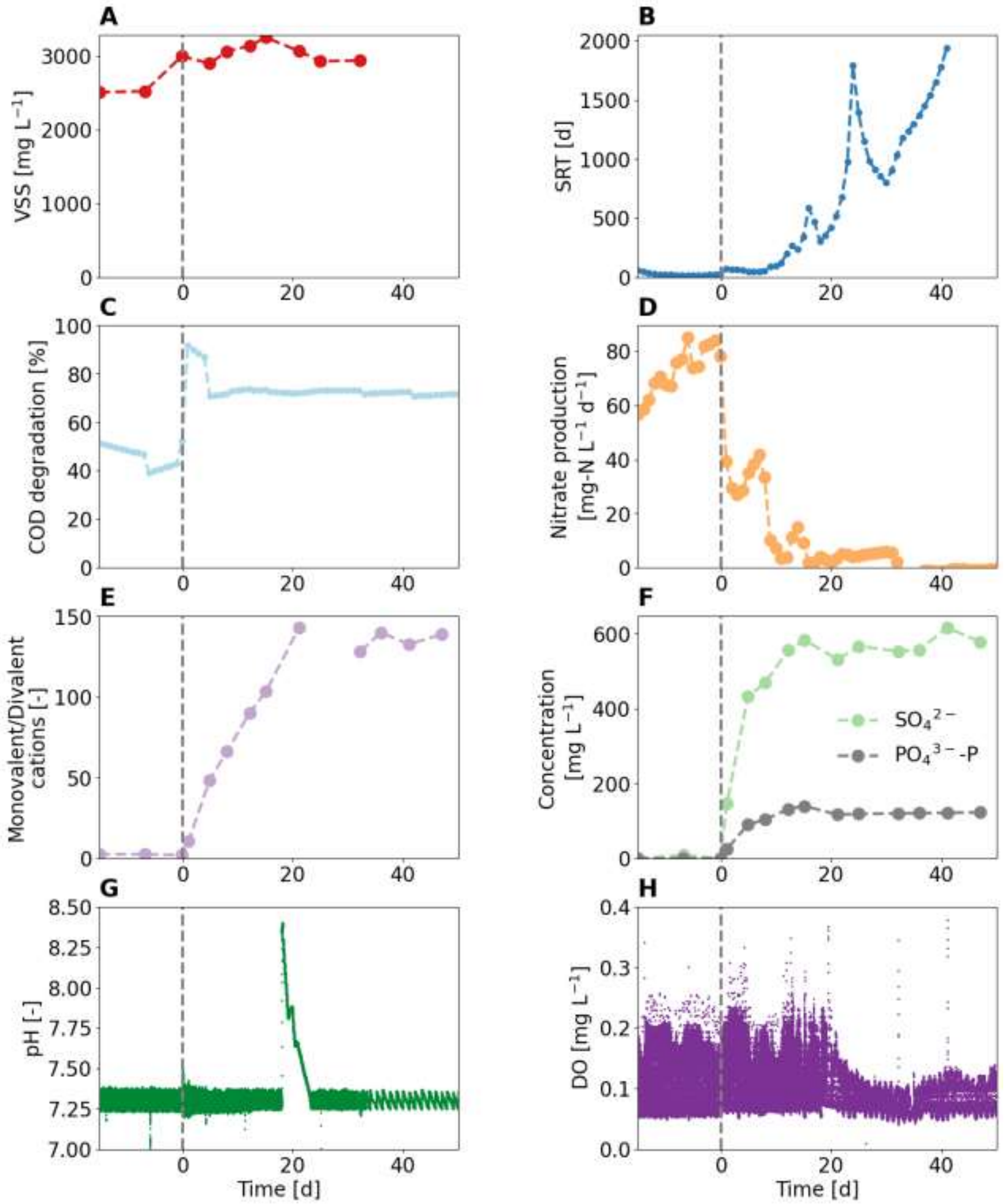

**Figure S6**: Reactor performance of the one-stage reactor fed with source-separated urine (A) VSS concentration in the reactor (B) solid retention time (SRT) (C) COD degradation comparing the influent and reactor COD (D) nitrate production rate (E) monovalent to divalent cations ratio in the reactor (F) the sulfate and phosphate concentrations in the reactor (G) the pH in the reactor (H) the dissolved oxygen (DO) concentration in the reactor.

**Figure S7** shows the microbial community composition in terms of AerAOB, NOB, and AnAOB.

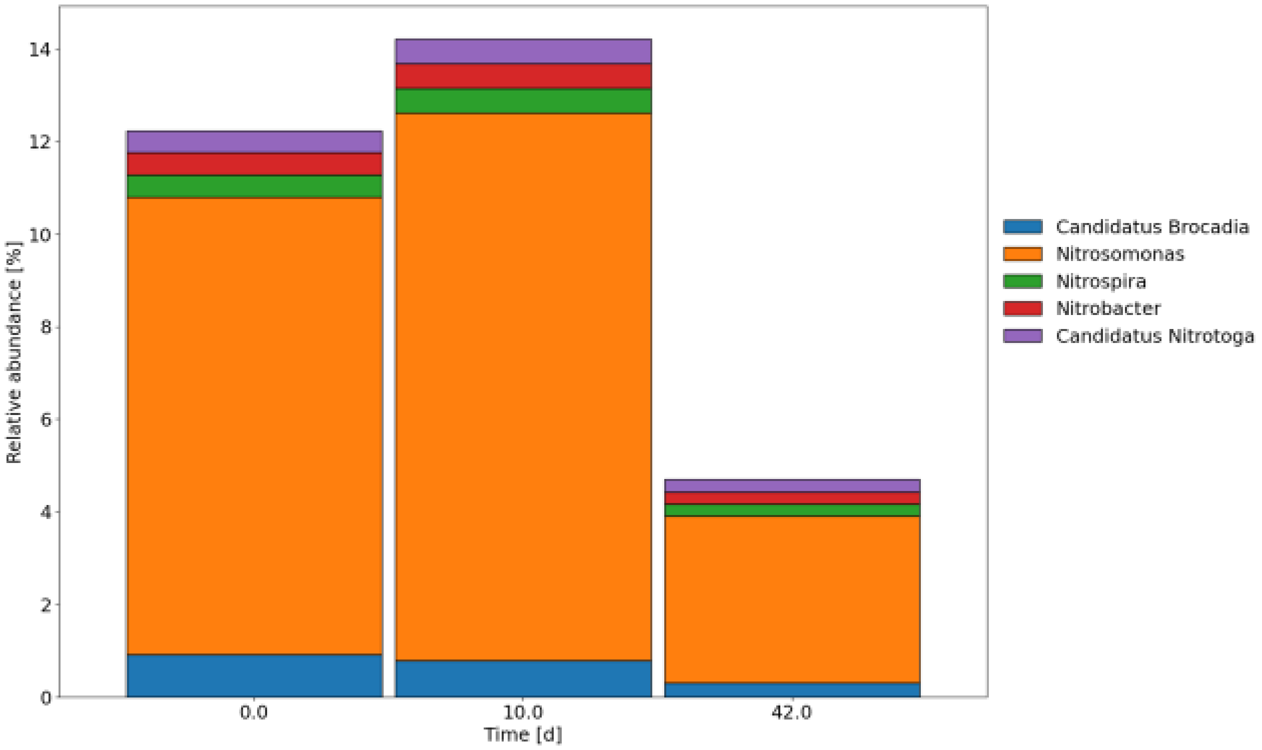

**Figure S7**: Relative read abundance of recognised genera of AerAOB, NOB, and AnAOB in the floccular sludge.

### 8 Reactor performance of one stage PN/A: synthetic urine

**Figure S8A** shows the VSS concentration in the reactor, **Figure S8B** the solid retention time (SRT), **Figure S8C** the COD degradation comparing the influent and reactor COD, **Figure S8D** the nitrate production rate, **Figure S8E** the monovalent to divalent cations ratio in the reactor, **Figure S8F** the sulfate and phosphate concentrations in the reactor, **Figure S8G** the pH in the reactor, **Figure S8H** the dissolved oxygen (DO) concentration in the reactor.

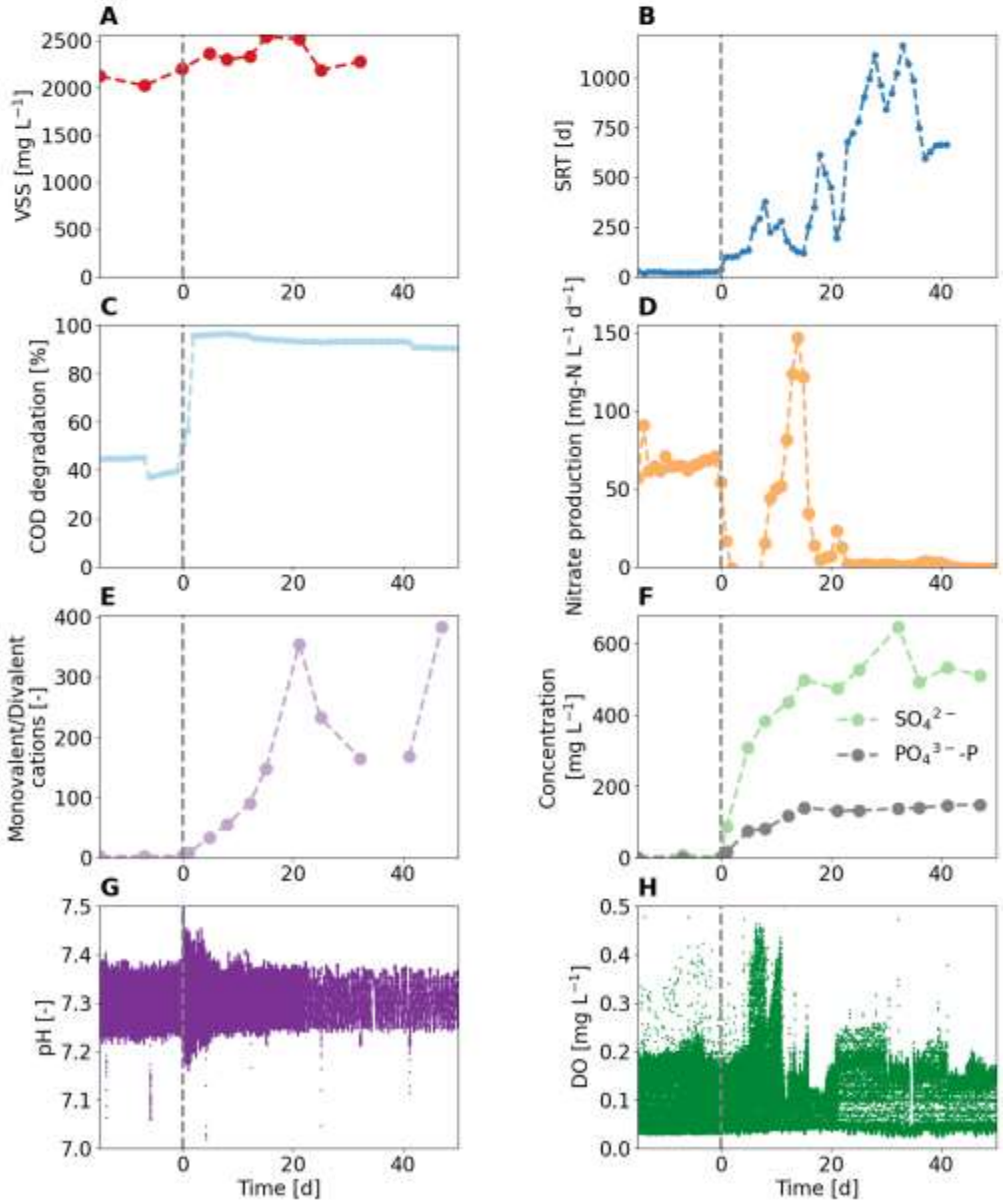

**Figure S8**: Reactor performance of the one-stage reactor fed with synthetic urine. (A) VSS concentration in the reactor (B) solid retention time (SRT) (C) COD degradation comparing the influent and reactor COD (D) nitrate production rate (E) monovalent to divalent cations ratio in the reactor (F) the sulfate and phosphate concentrations in the reactor (G) the pH in the reactor (H) the dissolved oxygen (DO) concentration in the reactor.

### 9 Reactor performance of one stage PN/A: source-separated urine with additional divalent cations (high concentration)

**Figure S9A** shows the VSS concentration in the reactor, **Figure S9B** the solid retention time (SRT), **Figure S9C** the COD degradation comparing the influent and reactor COD, **Figure S9D** the nitrate production rate, **Figure S9E** the monovalent to divalent cations ratio in the reactor, **Figure S9F** the sulfate and phosphate concentrations in the reactor, **Figure S9G** the pH in the reactor, **Figure S9H** the dissolved oxygen (DO) concentration in the reactor.

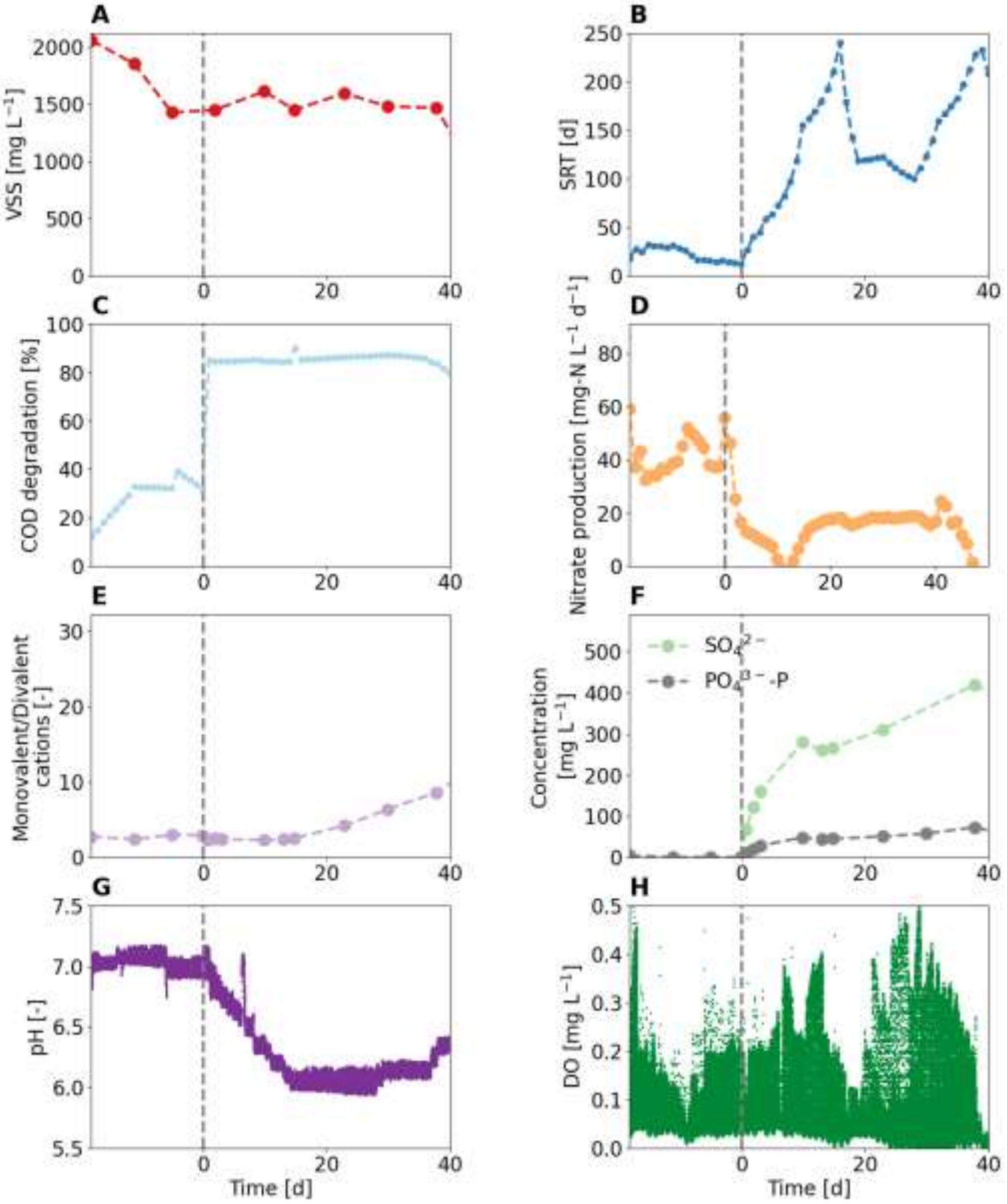

**Figure S9**: Reactor performance of the one-stage reactor fed with source-separated urine and additional divalent cations. (A) VSS concentration in the reactor (B) solid retention time (SRT) (C) COD degradation comparing the influent and reactor COD (D) nitrate production rate (E) monovalent to divalent cations ratio in the reactor (F) the sulfate and phosphate concentrations in the reactor (G) the pH in the reactor (H) the dissolved oxygen (DO) concentration in the reactor.

#### 10 Reactor performance of one stage PN/A: source-separated urine with biocarriers

**Figure S10A** shows the VSS concentration in the reactor, **Figure S10B** the solid retention time (SRT), **Figure S10C** the COD degradation comparing the influent and reactor COD, **Figure S10D** the nitrate production rate, **Figure S10E** the monovalent to divalent cations ratio in the reactor, **Figure S10F** the sulfate and phosphate concentrations in the reactor, **Figure S10G** the pH in the reactor, **Figure S10H** the dissolved oxygen (DO) concentration in the reactor.

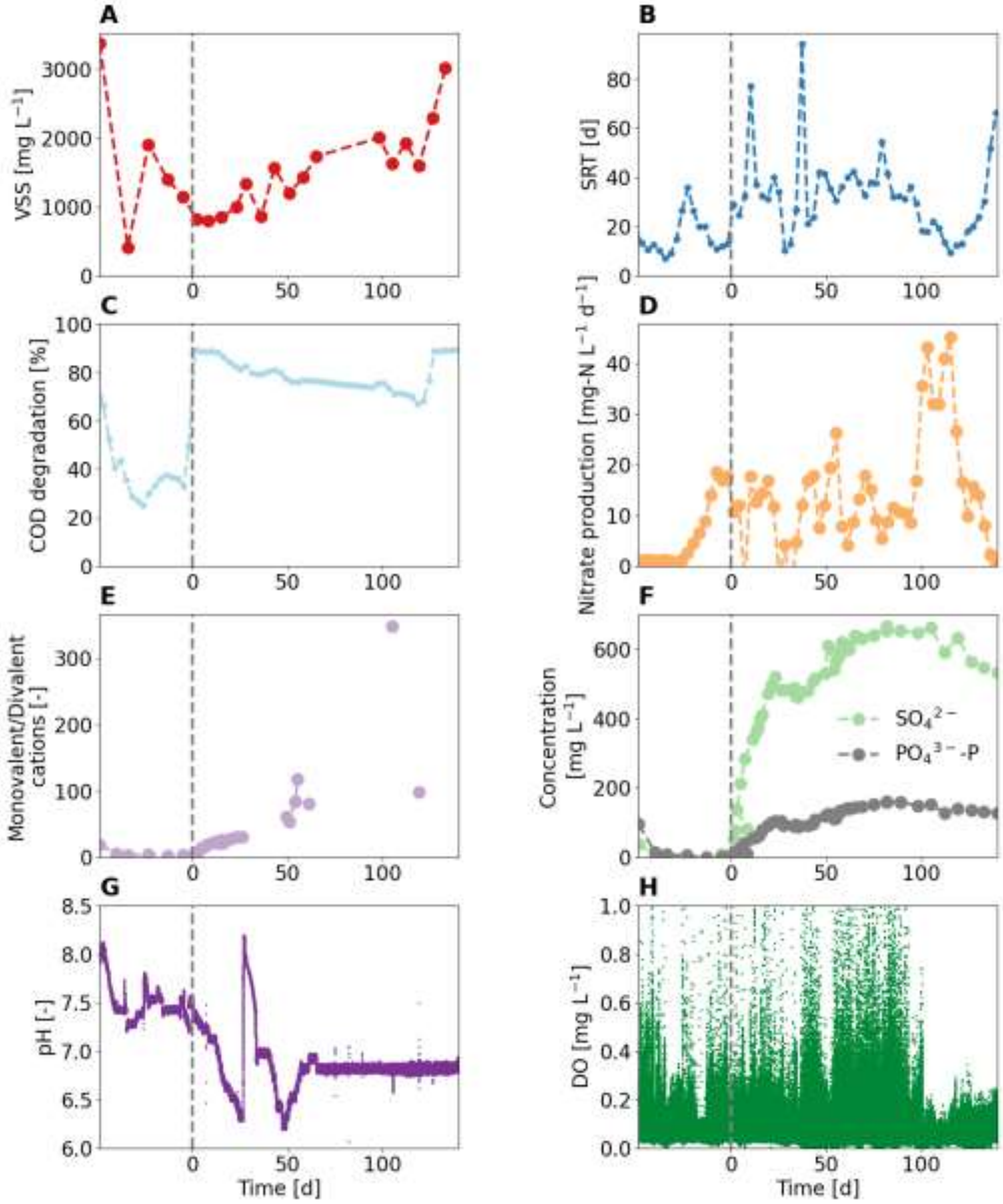

**Figure S10**: Reactor performance of the one-stage reactor fed with source-separated urine and bio-carriers. (A) VSS concentration in the reactor (B) solid retention time (SRT) (C) COD degradation comparing the influent and reactor COD (D) nitrate production rate (E) monovalent to divalent cations ratio in the reactor (F) the sulfate and phosphate concentrations in the reactor (G) the pH in the reactor (H) the dissolved oxygen (DO) concentration in the reactor.

### 11 Reactor performance of one stage PN/A: source-separated urine with additional divalent cations (low concentration)

Two experiments were carried out with additional divalent cations. As both failed, only the one with the higher concentration of divalent cations is included in the main manuscript. For completeness, the one with the lower amount of divalent cations is described here instead. **Table S11** and **Table S12** show the composition of the digester supernatant (start-up phase) and urine influent with the addition of divalent cations. The amount of calcium and magnesium added corresponds to the amount precipitated as calcium phosphate and struvite during urine storage. **Figure S11** shows the main result. The reactor failed after a short time. **Figure S12** shows the additional results.

**Table S11**: Digester supernatant used for the start-up of the additional experiment (n = 4).

| | | **Min** | **Mean** | **Max** | **Stdv.** |
|---|---|---|---|---|---|
| Chloride | [mg $L^{-1}$] | 139 | 139 | 140 | 0.5 |
| Phosphate | [mg-P $L^{-1}$] | <1 | <1 | <1 | NA |
| Sulfate | [mg $L^{-1}$] | <3 | <3 | <3 | NA |
| Sodium | [mg $L^{-1}$] | 71 | 74 | 75 | 2 |
| Ammonium | [mg-N $L^{-1}$] | 688 | 697 | 700 | 6 |
| Potassium | [mg $L^{-1}$] | 80 | 82 | 85 | 2 |
| Calcium | [mg $L^{-1}$] | 6 | 78 | 117 | 26 |
| Magnesium | [mg $L^{-1}$] | 42 | 42 | 43 | 0.5 |
| $COD_{sol}$ | [mg-$O_2$ $L^{-1}$] | 171 | 193 | 207 | 15 |
| Conductivity | [mS $cm^{-1}$] | 5.8 | 5.9 | 5.9 | 0.03 |
| pH | [-] | 8.1 | 8.2 | 8.2 | 0.03 |
| Temperature | [°C] | 18 | 19 | 20 | 0.8 |

**Table S12**: Source-separated urine with low concentration of additional divalent cations used for the additional experiment (n = 8). *estimated from reactor concentration

| | | **Min** | **Mean** | **Max** | **Stdv.** |
|---|---|---|---|---|---|
| Chloride | [mg $L^{-1}$] | 2070 | 2350 | 2660 | 20 |
| Phosphate | [mg-P $L^{-1}$] | 120 | 140 | 180 | 20 |
| Sulfate | [mg $L^{-1}$] | 490 | 542 | 626 | 47 |
| Sodium | [mg $L^{-1}$] | 830 | 1170 | 1440 | 170 |
| Ammonium | [mg-N $L^{-1}$] | 1470 | 2410 | 2790 | 470 |
| Potassium | [mg $L^{-1}$] | 770 | 1080 | 1230 | 140 |
| Calcium* | [mg $L^{-1}$] | | 34 | | |
| Magnesium* | [mg $L^{-1}$] | | 19 | | |
| $COD_{sol}$ | [mg-$O_2$ $L^{-1}$] | 770 | 990 | 1820 | 340 |
| Conductivity | [mS $cm^{-1}$] | 19.1 | 21.1 | 24.1 | 1.7 |
| pH | [-] | 8.76 | 8.85 | 8.88 | 0.05 |
| Temperature | [°C] | 18.2 | 20.1 | 21.7 | 1.4 |

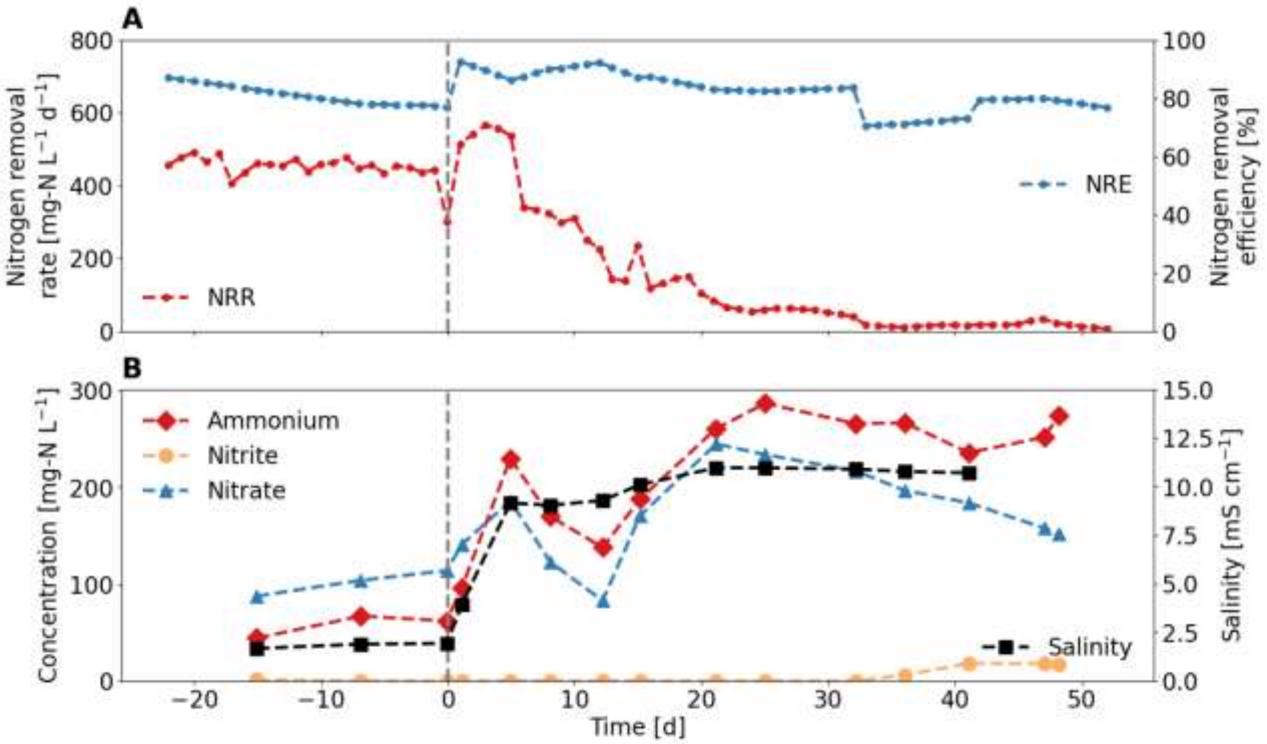

**Figure S11**: Reactor initially fed with digester supernatant and switched (dotted line) to source-separated urine with additional divalent cations. (A) Nitrogen removal rate (NRR) and nitrogen removal efficiency (NRE). (B) Nitrogen compounds and salinity measured as electric conductivity in the effluent.

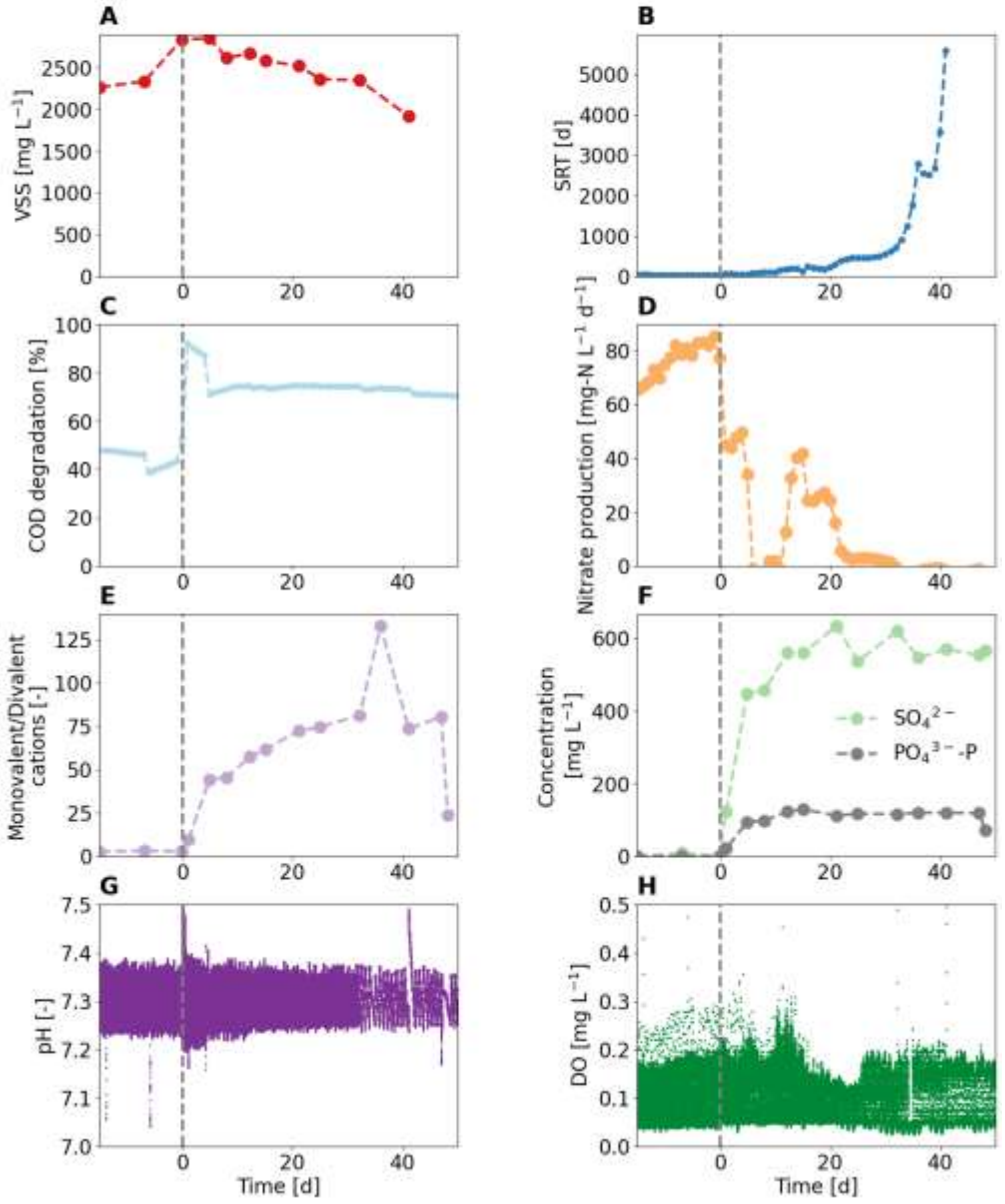

**Figure S12**: Reactor performance of the one-stage reactor fed with source-separated urine and bio-carriers. (A) VSS concentration in the reactor (B) solid retention time (SRT) (C) COD degradation comparing the influent and reactor COD (D) nitrate production rate (E) monovalent to divalent cations ratio in the reactor (F) the sulfate and phosphate concentrations in the reactor (G) the pH in the reactor (H) the dissolved oxygen (DO) concentration in the reactor.

### 12 Reactor performance of one stage PN/A: digester supernatant with additional salts

An additional experiment was carried out where salts (NaCl, KCl, and towards the end also sulfate) were added to the digester supernatant. As the NRR was continuously decreasing prior to the influent switch, no meaningful conclusions could be drawn from this experiment. Therefore, it was not included in the main manuscript. For completeness, it is described in detail here. **Table S13** and **Table S14** show the composition of the digester supernatant (start-up phase) and digester supernatant with the addition of salts. **Figure S13** shows the main result. The reactor failed after a short time. **Figure S14** shows the additional results.

**Table S13**: Digester supernatant used for the start-up of the additional experiment (n = 5).

| | | Min | Mean | Max | Stdv. |
|---|---|---|---|---|---|
| Chloride | [mg $L^{-1}$] | 144 | 151 | 160 | 6 |
| Phosphate | [mg-P $L^{-1}$] | <1 | <1 | <1 | NA |
| Sulfate | [mg $L^{-1}$] | <3 | <3 | <3 | NA |
| Sodium | [mg $L^{-1}$] | 78 | 87 | 102 | 9 |
| Ammonium | [mg-N $L^{-1}$] | 574 | 671 | 830 | 98 |
| Potassium | [mg $L^{-1}$] | 84 | 100 | 116 | 12 |
| Calcium | [mg $L^{-1}$] | 36 | 48 | 75 | 16 |
| Magnesium | [mg $L^{-1}$] | 38 | 42 | 50 | 5 |
| $COD_{sol}$ | [mg-$O_2$ $L^{-1}$] | 207 | 234 | 272 | 29 |
| Conductivity | [mS $cm^{-1}$] | 5.2 | 5.6 | 6.0 | 0.3 |
| pH | [-] | 8.13 | 8.25 | 8.23 | 0.09 |
| Temperature | [°C] | 21.0 | 22.0 | 22.9 | 0.7 |

**Table S14**: Digester supernatant with additional salts used for the additional experiment (n = 9). *estimated from reactor concentration. ** Sulfate was added to further increase salinity

| | | Min | Mean | Max | Stdv. |
|---|---|---|---|---|---|
| Chloride | [mg $L^{-1}$] | 1860 | 2220 | 2750 | 380 |
| Phosphate | [mg-P $L^{-1}$] | <1 | <1 | <1 | <1 |
| Sulfate** | [mg $L^{-1}$] | 10 | 290 | 744 | 349 |
| Sodium | [mg $L^{-1}$] | 900 | 1070 | 1340 | 190 |
| Ammonium | [mg-N $L^{-1}$] | 512 | 656 | 743 | 74 |
| Potassium | [mg $L^{-1}$] | 911 | 1066 | 1316 | 174 |
| Calcium* | [mg $L^{-1}$] | 47 | 60 | 70 | 12 |
| Magnesium* | [mg $L^{-1}$] | 7 | 23 | 44 | 16 |
| $COD_{sol}$ | [mg-$O_2$ $L^{-1}$] | 279 | 302 | 338 | 26 |
| Conductivity | [mS $cm^{-1}$] | 11.1 | 13.0 | 15.3 | 2.1 |
| pH | [-] | 8.1 | 8.2 | 8.4 | 0.1 |
| Temperature | [°C] | 24.4 | 25.7 | 28.7 | 1.8 |

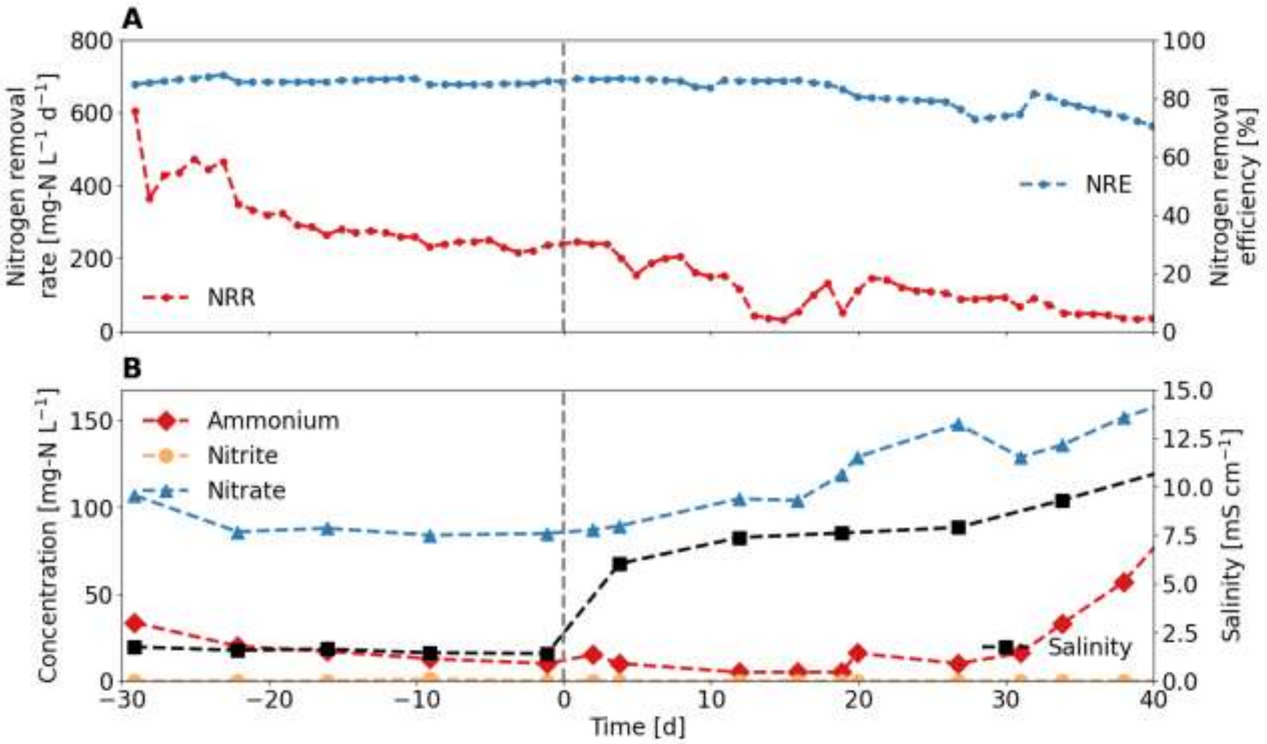

**Figure S13**: Reactor initially fed with digester supernatant and switched (dotted line) to digester supernatant with additional salts. (A) Nitrogen removal rate (NRR) and nitrogen removal efficiency (NRE). (B) Nitrogen compounds and salinity measured as electric conductivity in the effluent.

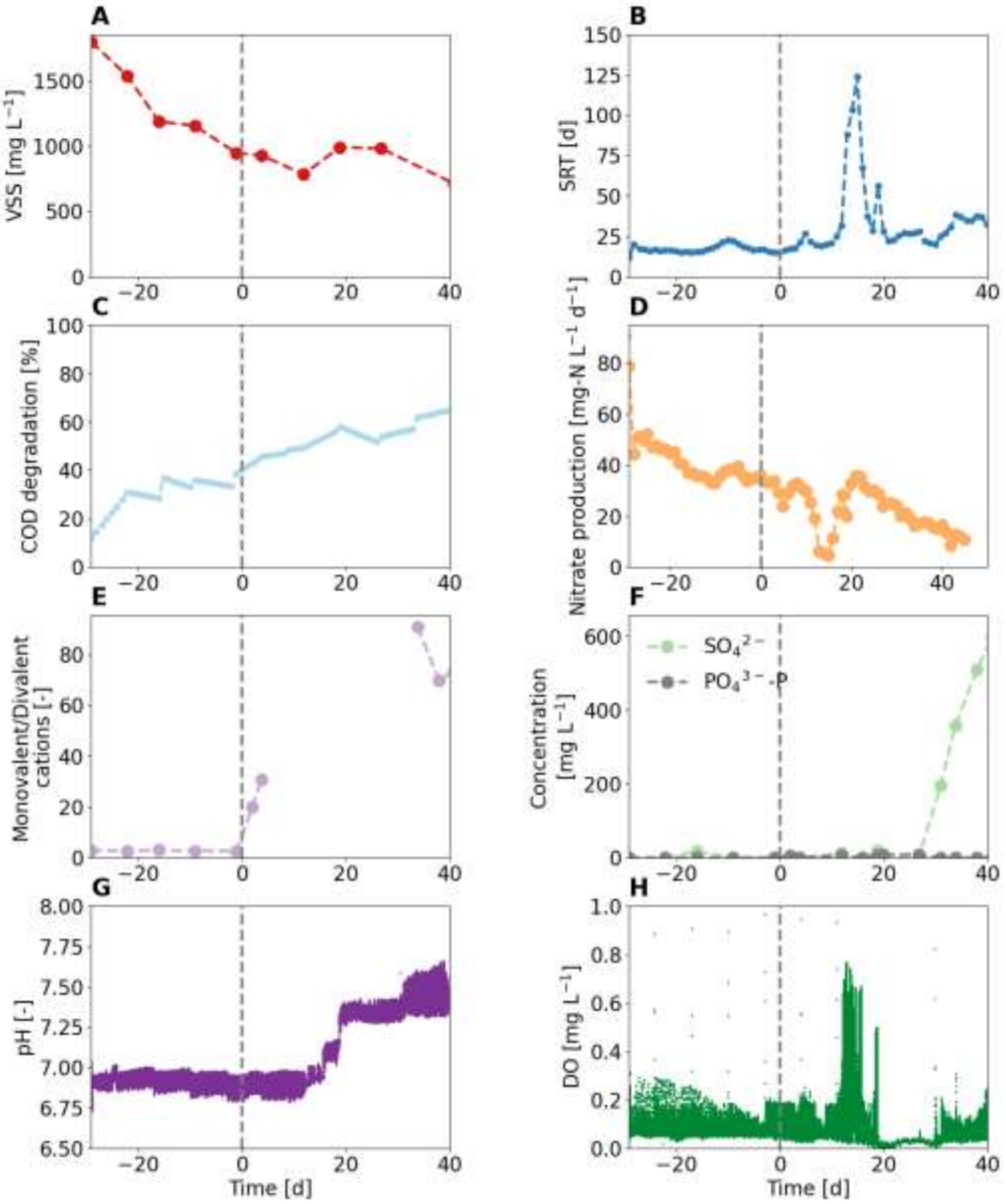

**Figure S14**: Reactor performance of the one-stage reactor fed with digester supernatant with additional salts. (A) VSS concentration in the reactor (B) solid retention time (SRT) (C) COD degradation comparing the influent and reactor COD (D) nitrate production rate (E) monovalent to divalent cations ratio in the reactor (F) the sulfate and phosphate concentrations in the reactor (G) the pH in the reactor (H) the dissolved oxygen (DO) concentration in the reactor.

#### 13 Ionic strength

**Figure S15** shows the ionic strength, calculated from the dissolved ion concentrations, plotted against the nitrogen removal rate (NRR).

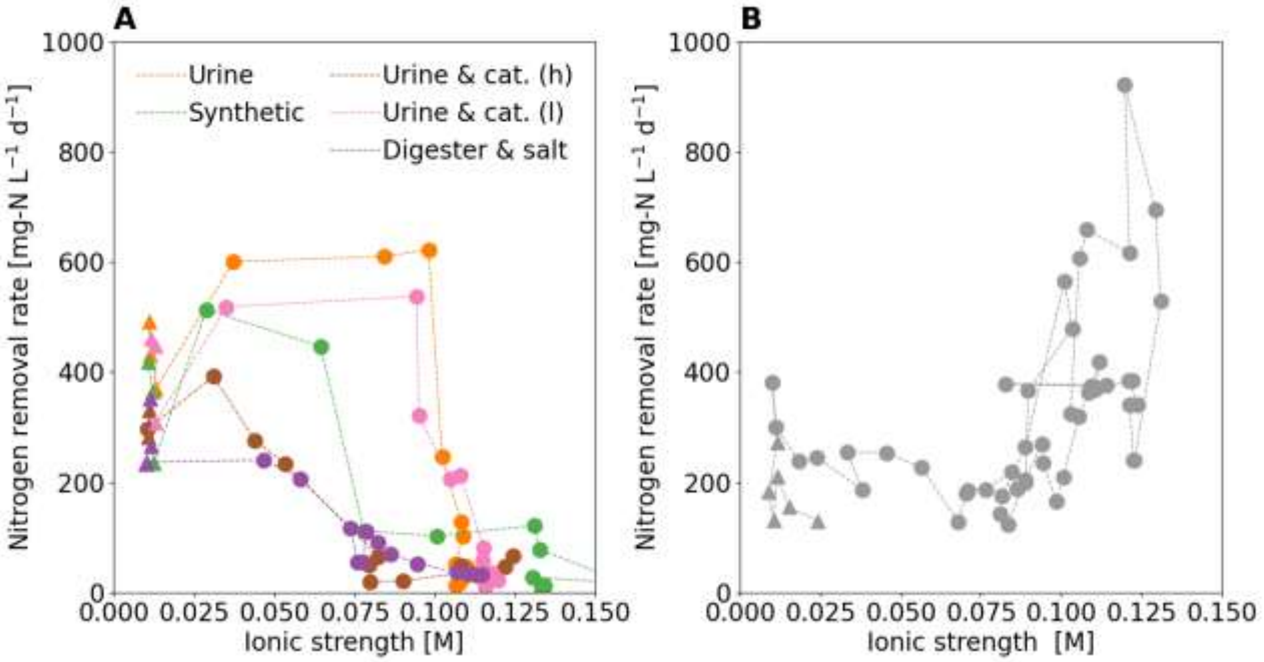

**Figure S15**: Ionic strength and nitrogen removal rate. Triangles (▲) represent the datapoints before the influent switch and circles (●) represent the data points after the influent switch. (A) In the reactors operated with floccular sludge, the nitrogen removal rate decreased with increasing ionic strength. (B) In the reactor operated with floccular sludge and biofilm carriers (hybrid), the nitrogen removal rate was maintained or even increased after the influent switch.
